# Gut community-level analysis reveals an altered balance between *Phocaeicola vulgatus* and *Bacteroides fragilis* in Alzheimer’s disease

**DOI:** 10.64898/2026.08.12.743985

**Authors:** Ziyuan Huang, Patrick M. McGrath, Danielle C. Ferdinand, Beth A. McCormick, Doyle V. Ward, Vanni Bucci, John P. Haran

**Author notes:** Corresponding author: Ziyuan Huang.

## Abstract

Gut microbiome differences in Alzheimer’s disease (AD) are typically cataloged taxon by taxon, yet bacterial competition and cross-feeding make species’ roles dependent on the entire community. We analyzed 274 stool metagenomes from 119 older adults (18 with AD) as communities, retaining 22 recurring across 1,000 runs. Using our AI framework, we identified 15 species differing in abundance in AD, particularly the commensal *Phocaeicola vulgatus* (Cohen’s *d* –0.91, 95% CI [-1.23, –0.59]), a finding robust to repeated sampling. It correlated negatively with its sister species, *Phocaeicola dorei* (*r* –0.57), suggesting possible niche competition; this replicated in an independent cohort (*r* –0.43). *P. vulgatus* was depleted in AD and the opportunistic pathogen *Bacteroides fragilis* enriched, shifting their balance toward *B. fragilis* (*d* –0.70), a modestly reproduced AD-associated pattern (*d* –0.24). Our findings suggest that AD-associated gut microbiome variation extends beyond taxon-specific abundance to the balance between specific species within a community matrix.

## INTRODUCTION

Alzheimer’s disease (AD) is a progressive neurodegenerative disorder and a leading cause of dementia among older adults. An estimated 7.2 million Americans aged 65 and older are living with AD, a number projected to reach 13.8 million by 2060 (1), and the caregiving burden that follows places growing demands on families, caregivers, and health systems. The microbiota-gut-brain axis has emerged as a promising avenue into that biology, linking the intestinal microbiome to neurodegenerative processes, yet the field remains in its early stages. Alterations in gut microbial composition have been associated with neuroinflammation in patients and mouse models (2–5), and individual taxa have been reported to activate microglia and worsen pathology (5). The available studies nonetheless remain heterogeneous in their cohorts, sampling schemes, and analytic design (6–10), so findings that appear in one cohort often fail to appear in the next. These differences make cross-study comparisons difficult and underscore the value of gut microbial associations with AD that are reproducible and externally validated.

Gut bacteria do not act alone. They form interdependent communities whose members compete for the same nutrients and cross-feed on one another’s products (11), so what a microbe does depends as much on its neighbors as on its own functions. Competition is sharpest where niches overlap. Close relatives usually co-occur across the human microbiome, and exclusion is typically found between distant relatives that share metabolic capacity (12), so competition tracks functional similarity more than ancestry. These interactions may help explain conflicting associations between gut bacterial taxa and AD. For example, Vogt et al. reported lower relative abundance of *Bifidobacterium* in individuals with AD (13), whereas Ling et al. observed higher abundance in another AD cohort (8). This discrepancy may reflect differences in study design, population, analytical methods, or species composition within the genus. We propose that a taxon association with AD may also depend on the surrounding community. A community therefore has structure that a single abundance cannot express, since one species can belong to several communities at once, and what a member’s depletion means may depend on which neighbor is abundant where that species is scarce. Yet most gut microbiome studies of AD still test each taxon on its own, returning catalogs of enriched and depleted species that vary from cohort to cohort. Testing species one at a time discards the covariation that organizes the community, and it cannot separate disease-specific differences from those that reflect gut ecology more broadly.

Community-level methods address that limitation directly. Latent Dirichlet Allocation (LDA) groups co-occurring bacteria into latent communities that can be analyzed as units (14, 15), and the approach has been applied to microbiome data alongside conventional differential abundance testing to give a fuller picture of the community (16, 17). However, the communities LDA recovers are unstable across random runs, so a community identified from a single model fit may not reappear in another run (18–20). A community read from one fit is therefore a hypothesis about structure rather than a finding. AD-microbiome associations, whether at the single-taxon or community level, are also seldom tested in a validation cohort, leaving their generalizability unresolved. Grouping species into communities therefore helps only if the communities themselves are reproducible and if the associations they carry are tested outside the cohort that produced them.

We addressed both requirements in the Gut-brain Alzheimer’s disease Inflammation and Neurocognitive Study (GAINS), a prospective cohort of community-dwelling older adults (21). We used our previously described Alzheimer’s disease Analysis Model (ADAM), a large language model framework that integrates microbiome and clinical data with evidence from the AD literature (22), to prioritize species for further investigation. Within this framework, we developed Meta-Topic Latent Dirichlet Allocation (MT-LDA), which identifies candidate communities across 1,000 model runs, retains those that recur, and estimates their associations with AD. We then carried the resulting species relationships, rather than the species list, into an independent amyloid-defined cohort to test whether the relationship holds. We hypothesized that reading gut microbiome as reproducible communities of co-occurring species, rather than analyzing species one at a time, would reveal AD-associated organization that single-species analyses miss. That approach brought two relationships within a single bacterial Family into view, one that holds with and without disease. We discovered a negative correlation, among AD patients and controls, between *Phocaeicola vulgatus* and *Phocaeicola dorei* that we externally validated. We also found an AD-associated difference in the balance between *P. vulgatus* and *B. fragilis*. These novel relationships emerged only after our ADAM-aided analysis of gut microbial community structure.

## RESULTS

### Individual species, not overall diversity or composition, differ in AD

We analyzed 274 stool metagenomes from 119 community-dwelling older adults in the GAINS cohort, 18 of whom had AD (47 AD samples and 227 control samples; **Fig. 1a** and **Extended Data Table 1**). The AD group differed in frailty, selective serotonin reuptake inhibitor use, and polypharmacy. All models were adjusted for the ten clinical characteristics in **Extended Data Table 1** plus visit day. Shannon diversity was only marginally lower in AD (adjusted difference –0.13, 95% CI [-0.38, 0.11]), and overall composition did not separate cases from controls (permutational multivariate analysis of variance on Bray-Curtis distance, PERMANOVA, *P* = 0.23); sex, not disease, structured the community (*P* = 0.001).

**Fig. 1.**
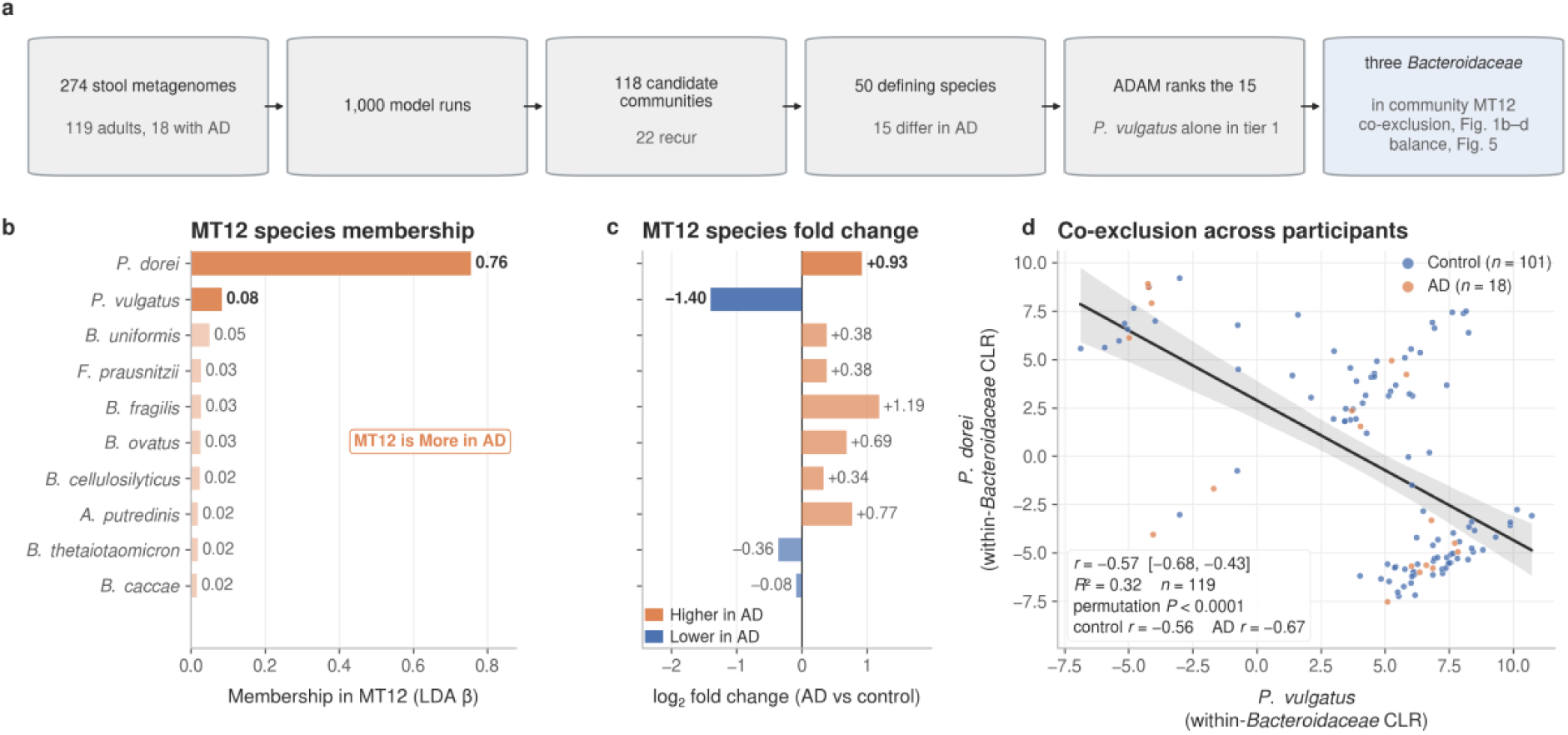
*Phocaeicola vulgatus* and its congener *Phocaeicola dorei* have negatively correlated abundances across participants. **a**, Study design and analysis workflow, from 274 stool metagenomes of 119 older adults, 18 with Alzheimer’s disease (AD), through community modeling across 1,000 runs to species prioritization using the **Alzheimer’s Disease Analysis Model (ADAM)**, which places *P. vulgatus* alone in tier 1. *P. dorei* has the highest membership weight in community MT12, which also includes *P. vulgatus* and *Bacteroides fragilis*. **b,** Mean membership weights for the ten leading species in MT12, representing each species’ contribution to the community across model runs. Orange bars indicate that MT12 is more abundant in AD, with the two *Phocaeicola* species listed at the top. **c,** Covariate-adjusted log_2_ fold changes in AD for the same ten species, ordered as in **b** orange and blue indicate higher and lower abundances in AD, respectively. **d,** *P. dorei* against *P. vulgatus*, shown as centered log-ratio (CLR) abundance within the 14 prevalent *Bacteroidaceae*, with one point per participant, blue for control and orange for AD. The line is the least-squares fit, and the band is its bootstrap 95% confidence interval over 1,000 resamples. Pearson r = −0.57, 95% CI [−0.68, −0.43] by Fisher z; R^2^ = 0.32; two-sided permutation P < 0.0001; r = −0.56 in controls and −0.67 in participants with AD. Panel **d** uses first-visit samples from 119 participants (101 controls and 18 with AD); panels **b** and c use 274 samples from the same participants.

Individual taxa told a different story. Of the 274 species tested, 23 were candidates, 18 higher in AD and 5 lower (**Extended Data Fig. 1**). The largest single difference was the enrichment of *Akkermansia* sp. KLE1605 (log_2_ fold change +1.42), followed by *Bacteroides fragilis* (+1.19), while the sharpest depletion was of *Phocaeicola vulgatus* (–1.40), the most abundant species in the cohort. Gut species do not act in isolation (11). A third of all species pairs co-varied in abundance across the 119 participants (12,468 of 37,401; median absolute Spearman correlation 0.34), and 695 conditional relationships among 260 species remained after adjusting each pair for all others (**Extended Data Fig. 2**). A species-by-species test discards this covariation, so we modeled the species as communities (**Fig. 1**).

### Twenty-two of 118 candidate communities recur across model runs

We grouped species into communities using latent Dirichlet allocation (LDA) (14), an unsupervised method that groups species whose abundances co-vary, and used LinDA (23), a linear model for compositional data, to estimate each community’s association with AD. LDA is a probabilistic model (14, 24); each run starts from a different random point and returns a similar but not identical set of communities. Four such runs produced different communities from different random starts and flagged different ones in each direction (**Extended Data Fig. 3**), so fitting the data once would tie the result to an arbitrary starting point.

We therefore ran LDA 1,000 times and pooled all 48,351 communities, 2,450 lower and 2,777 higher in AD, into a single volcano plot (**Fig. 2a**). Arranged by species composition rather than by AD effect, the communities that recurred across runs landed together as discrete islands, and the AD-leaning communities occupied a subset of these islands (**Fig. 2b**). We aligned the runs on this compositional similarity by nearest-match clustering against a consensus reference built by Hungarian assignment (25), and refer to each aligned community by a Meta-Topic (MT) code, a procedure we call MT-LDA (**Extended Data Fig. 4**).

**Fig. 2.**
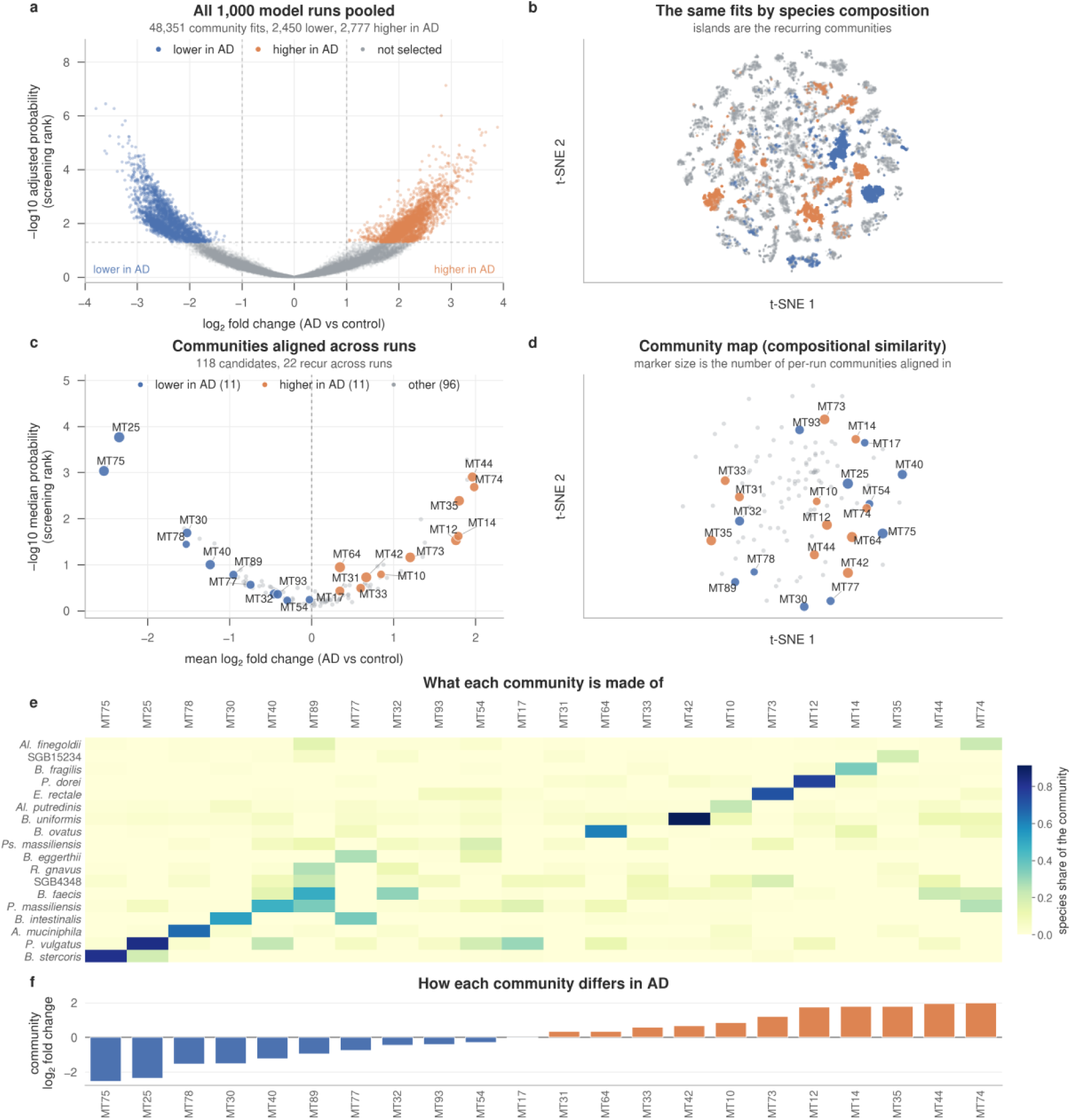
Twenty-two of 118 candidate communities recur across 1,000 model runs, 11 less abundant and 11 more abundant in Alzheimer’s disease. **a**, All 48,351 community fits from the 1,000 runs, plotted as log2 fold change in AD against –log10 Benjamini-Hochberg adjusted probability, read as a screening rank. Color marks the screen’s own selection: blue lower in AD, orange higher, gray not selected; 2,450 are lower and 2,777 higher. Dashed lines mark the screening thresholds: log2 fold change of 1 and adjusted probability of 0.05. **b**, The same fits placed by species composition. Both axes are unitless, and only local neighborhood structure is interpretable. Each island is a community the model recovered repeatedly from different starting points. **c**, The 118 candidate communities after alignment, colored where they meet the retention rule, 11 lower in AD and 11 higher, with the other 96 in gray and marker area scaling with the number of per-run communities aligned into each. **d**, The same 118 at their compositional centroids, identically colored and sized. **e**, Composition of the 22 recurring communities, each cell the mean weight a species carries in that community. **f**, The mean log_2_ fold change of each of the 22, in the same column order as **e**, spanning –2.53 in the *Bacteroides stercoris* community MT75 to +1.98 in MT74. Center values are covariate-adjusted fitted log_2_ fold changes over 274 samples from 119 participants, 18 with AD. No interval is drawn in **b** to **f**, and no probability beyond the screening ranks on the vertical axes of **a** and **c**.

Across the 1,000 runs, MT-LDA identified 22 recurring communities among 118 candidates (**Fig. 2c,d**), each recovered in a median of 766 runs and never fewer than 512, holding one AD direction in 77 to 100% of them. The 22 divided evenly, 11 lower in AD and 11 higher, a balanced split rather than a one-directional depletion (**Fig. 2f**). Community-level differences were large, spanning mean log2 fold changes from roughly –2.5 to +2.0, sharpest in the *Bacteroides stercoris* community MT75 (–2.53). Most communities were built around a distinct dominant species (**Fig. 2e**), and the lower– and higher-AD communities were compositionally intermixed rather than forming two separate clusters (**Fig. 2d**), so the AD difference is a rearrangement within the community structure rather than a split into two. Neither diversity nor overall composition captured this organization, which surfaced only when we analyzed the microbiome as communities.

### ADAM ranks *P. vulgatus* as the leading AD-depletion signal among complex modeling results

Fifteen of the 50 species that define the 22 communities differed in AD with intervals excluding zero, 10 lower and 5 higher, at effect sizes of roughly 0.3 to 0.9 (Cohen’s *d*; **Fig. 3a**). Several belonged to communities of opposite AD direction, so a species’ community memberships did not reduce to a single disease direction (**Fig. 3b,c**). *P. vulgatus*, for example, led three communities that were lower in AD (MT17, MT25, and MT54) yet was a prominent member of the higher-in-AD *Bacteroides ovatus* community MT64. No single effect size therefore summarizes a species’ relationship to AD, so ranking the 15 candidates required more than comparing their abundances. This is because ADAM aims to identify not only critical but also novel signals based on evidence from a particular study and literature analytics.

**Fig. 3.**
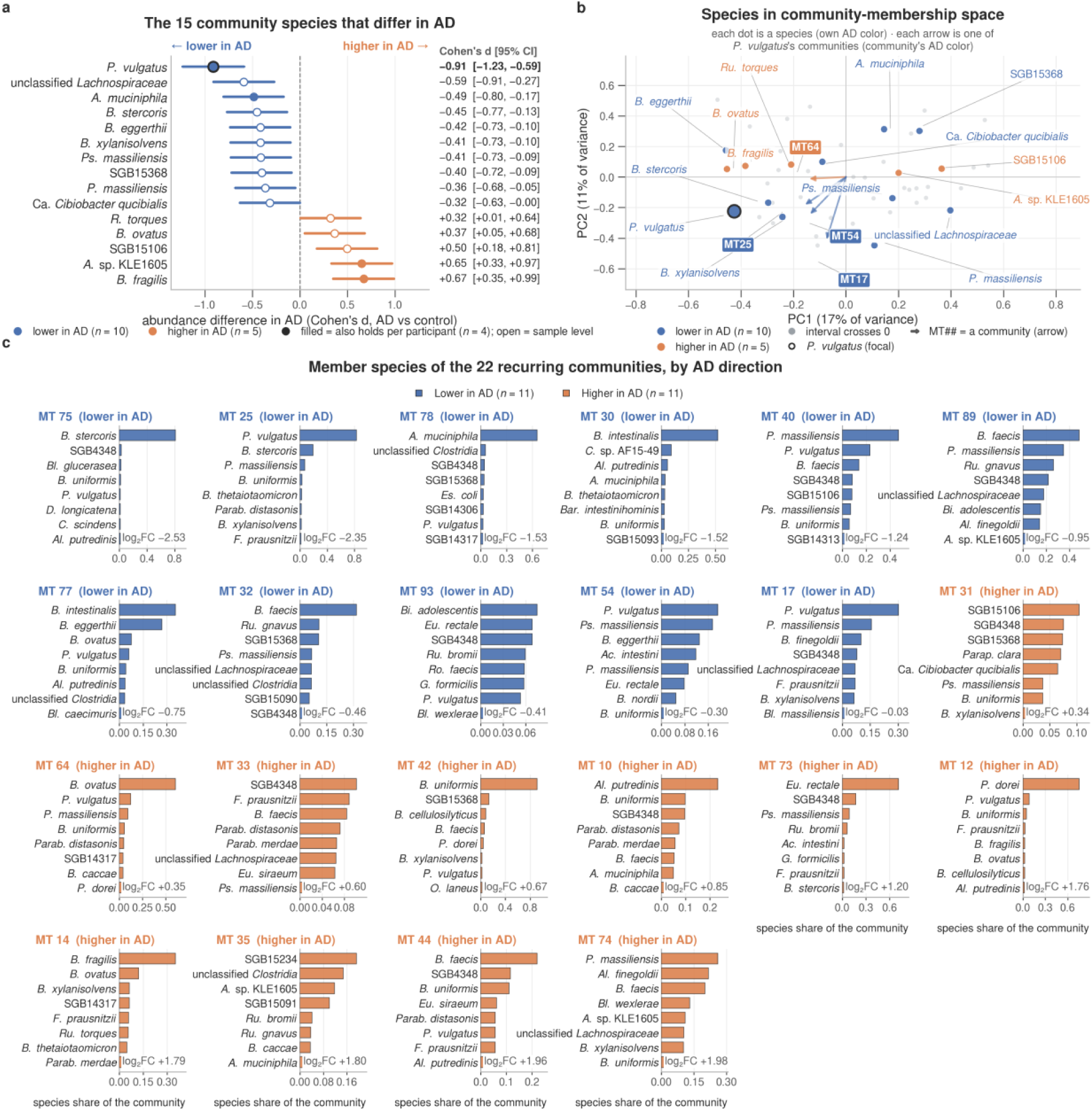
Fifteen of the 50 community species differ in Alzheimer’s disease, and they are spread across communities rather than gathered in one. **a**, The 15 species whose 95% confidence interval excludes zero, of the 50 that define the 22 recurring communities, ranked by effect. Dots show Cohen’s *d* for AD vs. control, and lines show its 95% confidence interval, each printed on the right. The range runs from *P. vulgatus* at –0.91, 95% CI [-1.23, –0.59], to *B. fragilis* at +0.67, 95% CI [+0.35, +0.99]. Blue is lower in AD (*n* = 10), and orange is higher (*n* = 5); a filled marker also holds at the participant level (*n* = 4). **b**, All 50 community species are placed by their community-membership profile. The 15 are colored by direction and named, and the remaining 35, whose intervals include zero, are gray. *P. vulgatus* carries a ringed marker. Four arrows run from the origin to the four communities the results discuss: MT17, MT25, and MT54, which *P. vulgatus* leads, and MT64, where it is the second member; each arrow matches its community’s direction color. The first component carries 17% of the variance and the second 11%. **c**, Each of the 22 recurring communities is shown as its own panel, ordered by that community’s mean log_2_ fold change from –2.53 in MT75 to +1.98 in MT74, with its eight highest-weight member species drawn as their weight within it. Eleven communities are lower in AD, and eleven are higher. Center values in **a** are Cohen’s *d* for AD against control on centered log-ratio abundance, with intervals from the Hedges-Olkin standard error, over 274 samples from 119 participants, 18 with AD. The community effects in **c** are covariate-adjusted fitted log_2_ fold changes.

The co-abundance network offered no shortcut, since the AD species were scattered throughout it and no more central or connected than others (**Extended Data Fig. 2**). We therefore applied our ADAM framework (22) to prioritize candidate species using six predefined criteria (**Table 1** and **Extended Data Fig. 8**). Only *P. vulgatus* reached tier 1. Its depletion is the largest of any species here (*d* = –0.91) and reliable, since it remains when each participant is counted once; it belongs to 9 of the 22 communities and leads three; and as an abundant, well-characterized plant-fiber fermenter (26, 27) that has not been extensively studied in AD, it is a fresh candidate for mechanistic work. Three species formed tier 2, *B. fragilis* and *Akkermansia muciniphila*, which lead communities with reliable differences but are already heavily reported in AD, and *Bacteroides stercoris*, which leads MT75 with little prior attention in AD, the closest second candidate. The remaining 11 formed tier 3, none combining community leadership with a strong, reliable, and focused difference; the enriched *Akkermansia* sp. KLE1605, the largest fold change in the species screen (**Extended Data Fig. 1**), finds no support in the community structure.

**Table 1.**
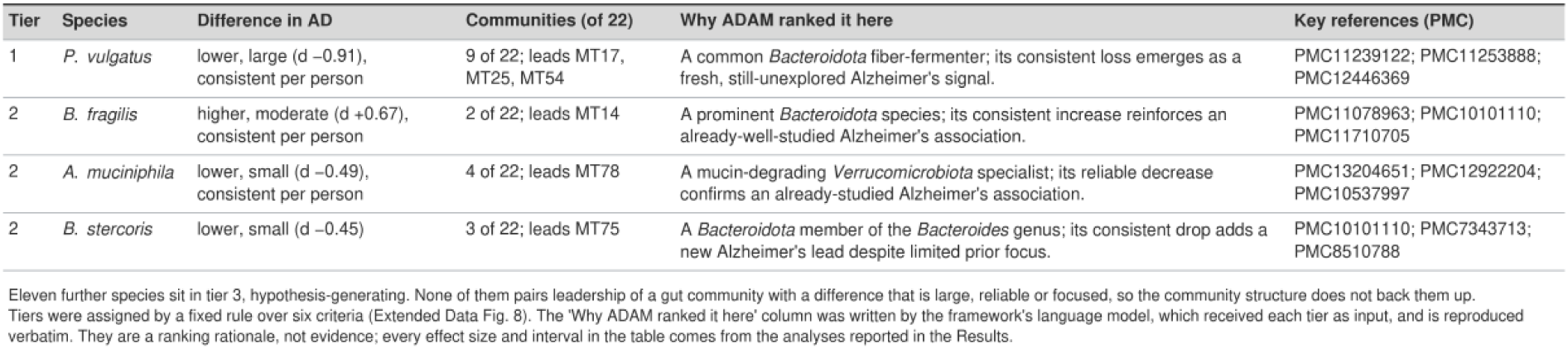
ADAM’s ranking of gut species as Alzheimer’s signals. The four species the ADAM framework places in tiers 1 and 2 are drawn from the 15 community species that differ in AD. Effect sizes are Cohen’s *d* for AD against control over 274 samples from 119 participants, 18 with AD, and each is a sample-level estimate whose 95% confidence interval excludes zero; the intervals themselves are drawn in **Fig. 3a**. “Consistent per person” marks a species whose participant-level interval also excludes zero, which 4 of the 50 community species do. A species enters that set of 50 by reaching 5% mean membership weight in at least one recurring community. The “Communities (of 22)” column counts a species as a member at 2% membership weight or more, with recurrence in at least half the model runs, a wider rule. Eleven further species sit in tier 3, hypothesis-generating, and none of them pair leadership of a gut community with a difference that is large, reliable, or focused. Tiers were assigned by a fixed rule with fixed thresholds over six criteria, five from the GAINS measurements and one from the count of articles co-mentioning the species and AD (**Extended Data Fig. 8**); no language model took part in the assignment. The “Why ADAM ranked it here” column was written by the **ADAM framework’s language model** from the assigned tier, the GAINS measurements, and the cited literature, and is reproduced as written. These are ranking rationales, not evidence, and every effect size in the table comes from the analyses reported in the results. The “Key references (PMC)” column gives PubMed Central identifiers retrieved by ADAM’s literature analytics layer.

A classical indicator species analysis (28) across all 274 species points to the same two Bacteroidaceae. *P. vulgatus* holds one of the highest indicator values (group-size-corrected indicator value 0.76, rank 3; permutation *P* = 0.028), earned as a common commensal present in 89% of participants without AD rather than as a specialist, whereas *B. fragilis* reaches its value through specificity instead (**Fig. 4a,b**). Taken as a whole, however, the screen returns 14 probabilities below 0.05, whereas 13.7 are expected by chance, so it describes the cohort rather than naming a biomarker.

**Fig. 4.**
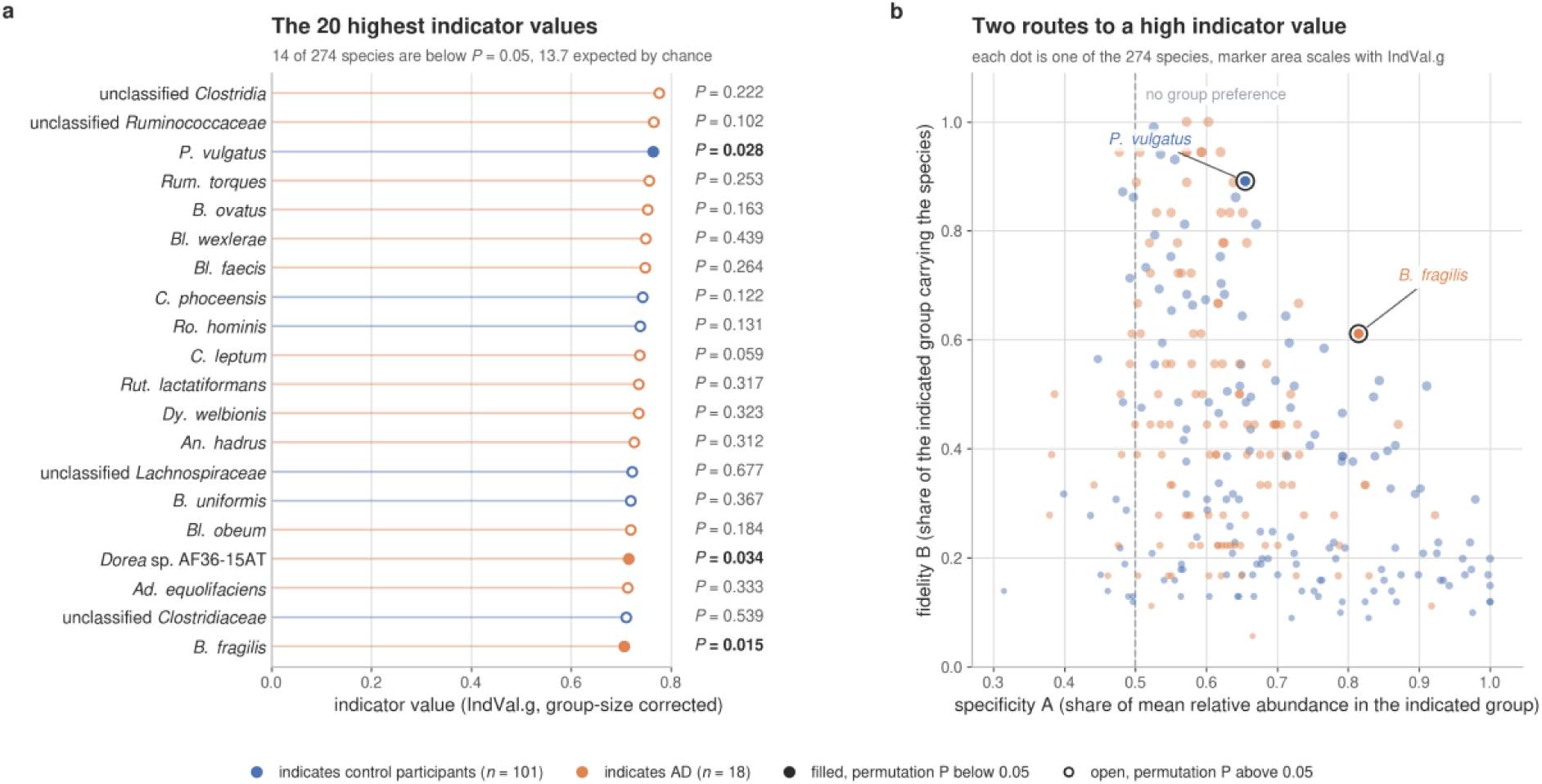
*Phocaeicola vulgatus* and *Bacteroides fragilis* lead the indicator species of Alzheimer’s disease status, at a hit count no larger than chance. Indicator species analysis over 274 species in 119 first-visit participants, 101 controls and 18 AD, by group-size-corrected indicator value with 9,999 permutations. **a**, The 20 highest indicator values. Blue marks a species that indicates control participants and orange one that indicates AD; a filled marker indicates a permutation probability below 0.05 and an open marker one above it, and each permutation probability is printed beside its species. Three of the 20 are below 0.05, *B. fragilis* at *P* = 0.015, *P. vulgatus* at *P* = 0.028, and *Dorea* sp. AF36-15AT at *P* = 0.034. *P. vulgatus* carries the highest indicator value of any species in the cohort below 0.05, 0.764, and *B. fragilis* has the highest specificity of the 20, 0.815. **b**, Specificity against fidelity for all 274 species, marker area scaling with the indicator value, with the dashed line at the specificity of no group preference. *P. vulgatus* and *B. fragilis* are ringed, and the two reach a high indicator value by different routes, *P. vulgatus* on fidelity and *B. fragilis* on specificity. Across all 274 species, 14 are below 0.05, and none survive the Benjamini-Hochberg adjustment. Therefore, these probabilities describe the cohort and do not identify any biomarker.

### *P. vulgatus* and *P. dorei* show co-exclusion across participants

ADAM’s integrative ranking singled out P. vulgatus, whose closest relative, P. dorei, shares community MT12. We examined how the leading signal relates to its congener. Within MT12, the two never co-dominate, *P. dorei* leading at a mean membership weight of 0.76 and *P. vulgatus* contributing a much smaller share (0.08; **Fig. 1b**). Most members of MT12 were higher in AD, whereas *P. vulgatus* was lower (log_2_ fold change –1.40) and *P. dorei* higher (+0.93; **Fig. 1c**). Examined directly, their within-*Bacteroidaceae* centered log-ratio (CLR) abundances (computed across the 14 prevalent *Bacteroidaceae* species) are strongly and negatively correlated across the 119 first-visit participants (*r* = –0.57, 95% CI [-0.68, –0.43]; permutation *P* < 0.0001), so a higher abundance of one congener coincides with a lower abundance of the other (**Fig. 1d**). We call this pattern co-exclusion (12), the inverse relationship being the observation and competitive exclusion the inference it invites. The negative correlation is present in both control and AD participants, so within GAINS the co-exclusion holds with and without disease.

### The P. vulgatus-to-B. fragilis balance favors B. fragilis in AD

A second signal within the *Bacteroidaceae* is *B. fragilis*, a gut opportunistic pathogen (29, 30) that is the most enriched of the 15 species that differ in AD (**Fig. 3a**) and the most enriched member of MT12 (log_2_ fold change +1.19; **Fig. 1c**). Its relationship to *P. vulgatus* is not a co-abundance, since the two are not coupled within participants, but a compositional balance (31), the log-ratio of the depleted commensal to the enriched opportunistic pathogen. Together with *P. dorei*, the three species make up about 13% of the gut community in control participants and 11% in AD, so the guild is similar in size in both groups, but its internal composition is not (**Fig. 5a**). *P. vulgatus* is about half as abundant in AD as in controls (5.9% against 11.0% of the community), *B. fragilis* is about four and a half times as abundant (1.45% against 0.32%), and *P. dorei* is about 70% more abundant (3.7% against 2.2%), so both neighbors are more abundant where *P. vulgatus* is scarce. Reduced to a single value per first-visit participant (n = 119), the balance is lower in AD (Cohen’s *d* = –0.70, 95% CI [-1.41, –0.10] in the whole-community frame; label-free permutation *P* = 0.009; **Fig. 5b**), and participants with AD sit toward the low-*P. vulgatus*, high-*B. fragilis* corner of the plane (**Fig. 5c**). The two relationships therefore differ. One is co-exclusion, present with or without disease, and the other is a balance that differs in AD, consistent with a balance that favors the opportunist over the commensal in AD. Whether either holds beyond GAINS is the question we take up next.

**Fig. 5.**
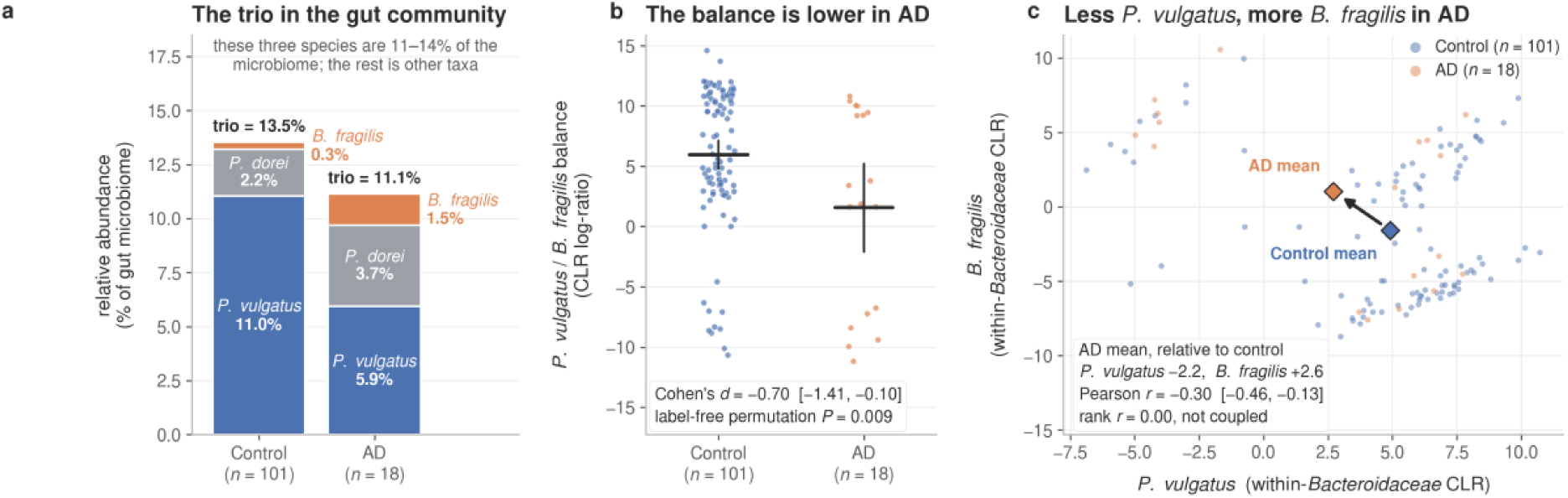
The *Phocaeicola vulgatus* to *Bacteroides fragilis* balance is lower in Alzheimer’s disease. **a**, Mean relative abundance of *P. vulgatus* (blue), *Phocaeicola dorei* (gray), and *B. fragilis* (orange), three members of the family *Bacteroidaceae* in community MT12 (Fig. 1b), in control participants and in participants with Alzheimer’s disease (AD). Their combined mean relative abundance is 13.5% in controls and 11.1% in participants with AD. **b,** The *P. vulgatus* to *B. fragilis* balance, defined as the whole-community centered log-ratio (CLR) abundance of *P. vulgatus* minus that of B. fragilis, one point per participant, blue for control and orange for AD. Horizontal bars mark group means, and vertical bars mark their bootstrap 95% confidence intervals over 2,000 resamples. Cohen’s d = −0.70, 95% CI [−1.41, −0.10] by bootstrap, two-sided permutation P = 0.009. **c,** *B. fragilis* against *P. vulgatus* using the within-*Bacteroidaceae* CLR transformation in Fig. 1d, with points colored as in **b**. Diamonds mark the group means, and the arrow connects the control mean to the AD mean. The AD mean is 2.2 CLR units lower for *P. vulgatus* and 2.6 units higher for *B. fragilis*. Pearson r = −0.30, 95% CI [−0.46, −0.13] by Fisher z; the rank correlation is 0.00. *B. fragilis* was not detected in approximately half of the participants. All panels use first-visit samples from 119 participants (101 controls and 18 with AD).

### Co-exclusion replicates in an independent cohort, the balance repeats direction

We tested the two GAINS relationships in AlzBiom, an independent AD case-control cohort (n = 175), beginning with the *P. vulgatus* and *P. dorei* co-exclusion. Applied unchanged, the co-exclusion reproduced (*r* = –0.43, 95% CI [-0.54, –0.30]; permutation *P* < 0.0001), a strength within 0.14 of GAINS (*r* = –0.57), with overlapping intervals (**Extended Data Fig. 5a,b**). The relationship was present in both control and AD participants across both cohorts (GAINS control *r* = –0.56, AD *r* = –0.67; AlzBiom control *r* = –0.49, AD *r* = –0.35). The *P. vulgatus* / *B. fragilis* balance is also lower in AD in AlzBiom (Cohen’s *d* = –0.24, 95% CI [-0.55, +0.05]), in the same direction as GAINS (*d* = –0.70) but attenuated, and its interval includes zero (**Extended Data Fig. 5c,d**). Of the two species-level differences that compose the balance, *B. fragilis* enrichment largely reproduces in AlzBiom whereas *P. vulgatus* depletion does not, so the balance rather than either species alone is what carries across. Both relationships hold under the within-*Bacteroidaceae* and the whole-community reference frames (**Extended Data Table 2**). The external test thus separates the two. The co-exclusion reproduces robustly and holds regardless of disease, a conserved property of the *Bacteroidaceae*, whereas the balance repeats only the guild’s AD-associated skew toward *B. fragilis*. Within the family, then, the disease-associated difference lies in the balance between two family members rather than in the depletion of one species alone.

## DISCUSSION

Our study reframes where the gut microbiome’s association with AD likely resides. Alongside the taxa that differential-abundance testing flagged, the signal in GAINS was organized among co-occurring species, so a taxon’s relationship to disease depended on the surrounding community. A single model run gives one arrangement of species and the next a different one; twenty-two of ours recurred across 1,000 runs and can be treated as units rather than artifacts of one fit, and that recurrence exposed a pattern single-species testing cannot show. Many species belonged to more than one community, and those communities did not all lean the same way in AD, so a coherent disease direction belonged to the community rather than to the species alone. *P. vulgatus* illustrates the distinction, consistently lower in AD yet a prominent member of the higher-in-AD *B. ovatus* community MT64. Species– and genus-level reports of *P. vulgatus* in AD and in older adults at risk of cognitive decline do not agree (32–38), which may reflect the same effect. As a fiber-fermenting commensal producing propionate, it is generally regarded as barrier-supporting and anti-inflammatory (39), whereas Gram-negative *Bacteroides* taxa are also discussed as contributors to lipopolysaccharide translocation, intestinal permeability, and microglial activation in AD (40). The same organism can therefore read as protective or as a marker of a harmful inflammatory context, depending on strain and community, and single-species testing captures only its depletion.

ADAM helped us interpret the community analyses by placing species’ abundance differences, consistency in participant-level analyses and community roles in the context of published AD evidence. Its language model generated biological explanations for priorities established by predefined criteria, with *P. vulgatus* along in the top tier (**Table 1 and Extended Fig. 8**). Its value lies in organizing complex results into a ranked shortlist with explicit rationales that can be checked against the underlying analyses and cited studies. Although co-mention counts remain an imperfect proxy for biological relevance, external validation covered only the pair it prioritized, so ADAM contextualizes and prioritizes rather than confirming. Support beyond ADAM’s ranking came from descriptive agreement with our indicator species analysis (28), in which *P. vulgatus* had the highest indicator value among species meeting the nominal significance threshold.

*P. vulgatus* and *P. dorei* are close congeners of the genus *Phocaeicola*, formerly *Bacteroides* (41), sharing roughly 96% sequence similarity (42) and the machinery for the same dietary and host glycans (27, 41), which predicts competition (11), and here the two do not both dominate the same gut; their abundances trading off rather than co-varying. The trade-off persists across both AD cohorts (**Extended Data Table 2**), the two diverge in inflammatory bowel disease (43) and in treatment-naive tuberculosis, where *P. dorei* is enriched in patients and *P. vulgatus* in controls (42), so although that evidence corroborates rather than replicates and the exclusion is not universal, it reads as a conserved constraint of gut ecology rather than an AD-specific artifact. The AD-associated depletion of *P. vulgatus* is therefore a perturbation of a conserved relationship, and which neighbor is more abundant in its place determines whether it matters to the host. A swap toward the close relative *P. dorei* is disease-neutral, present regardless of AD, although the two differ in carbohydrate utilization (41). The consequential difference involves a different neighbor, the aerotolerant opportunist *B. fragilis* (44), which colonizes the intestinal mucus layer (45) and is more abundant where *P. vulgatus* is scarce. Its relationship with the host is strain-contingent (46), nontoxigenic strains suppressing inflammatory signaling (45, 47) and enterotoxigenic strains carrying the fragilysin locus that cleaves epithelial junctions and converts a commensal into a colitogenic pathobiont (48). In mice, isolates from AD gut microbiomes activate microglia and trigger pathology in neuronal C/EBPβ transgenic animals (5) and *B. fragilis* raised plaque burden and suppressed microglial clearance of amyloid-β (49); neither establishes the phenotype of the strains more abundant in patients, but both show that *B. fragilis* can engage mucosal immunity in a way *P. dorei* cannot. A guild skewed from a luminal commensal toward a mucosa-associated opportunist alters which organism contacts the host rather than what the community produces, so any consequence for the brain would plausibly run through mucosal immunity and barrier integrity (40, 50, 51). We advance this only as a hypothesis, since relative enrichment of *B. fragilis* may reflect an altered gut environment as much as drive one, and the link to neuroinflammation is untested.

### Limitations

The primary limitation is the sample size of 18 participants with AD. Therefore, we report effect sizes with confidence intervals, and a null result indicates a lack of evidence rather than proof of no effect. Participants contributed repeated samples, so sample-level fits are population-averaged, not participant-level. By design, the whole pipeline is a feature-selection mechanism rather than a hypothesis test, so its consistency counts mark selection rather than confirmation. The estimates are in-sample and AD-label-selected, an optimistic bracket rather than out-of-sample inference. We tested the candidates against AlzBiom (52), which defines AD by probable dementia with cerebrospinal-fluid amyloid, a different biomarker route from GAINS, and against the literature through ADAM’s 174,230-article PMC corpus, which can mark a signal as candidate-novel but cannot validate it. Neither source confirms these candidates, so the next step is direct testing in experimental models.

### The AD signal resides in community composition, not in any one organism

These relationships emerge when the community is resolved to individual species. The congeners’ trade-off and the skew toward *B. fragilis* play out among species that a genus-level would merge into one *Bacteroidaceae* signal. Shotgun metagenomics resolves taxa to species (53), separating a conserved competitive relationship from a disease-associated one that a genus would average out, and community models read the same species in each neighborhood it occupies. Two relationships were carried to an independent cohort, a conserved co-exclusion that reproduced and held regardless of disease, and a disease-associated skew toward an opportunist within the same guild, whose direction repeated. Together, they show that AD-associated variation extends beyond taxon-specific abundance to a community’s composition. The result is a short set of externally grounded relationships rather than a longer list of taxa, each triangulated against the published record by ADAM. These exploratory, hypothesis-generating associations distill a complex community into a small, interpretable set of *Bacteroidaceae* relationships for larger prospective AD cohorts to confirm; the deeper frontier is the strain level, where the immunological character of *B. fragilis* is determined (45) and where the next test of this ecology must be conducted.

## METHODS

### Cohort and study design

We analyzed data from the GAINS cohort (21) with Alzheimer’s disease status as the primary outcome. We defined cohort groups within the National Institute on Aging and Alzheimer’s Association (NIA-AA) **“**ATN**”** research framework (54) using blood-based measures of amyloid and tau categories, the plasma Aβ42/Aβ40 peptide ratio, and phosphorylated tau217. As part of this experimental design for differential abundance analysis, we categorized participants as having Alzheimer’s disease or not (controls). The cohort comprises community-dwelling adults aged 60 years and older, recruited prospectively at the University of Massachusetts Chan Medical School.

### Ethics

The Gut-brain Alzheimer’s disease Inflammation and Neurocognitive Study (GAINS) was approved by the Institutional Review Board of the University of Massachusetts Chan Medical School (IRB docket H00021745) and conducted in accordance with the Declaration of Helsinki. Written informed consent was obtained from all participants or their legal representatives. Stool shotgun metagenomic sequencing and all data analyses were performed in-house. We describe the cohort design and clinical assessment protocol in our prior report (21).

### Metagenomic sequencing and taxonomic profiling

We collected stool samples from participants and extracted DNA using the DNeasy PowerSoil Pro Kit (Qiagen). The sequencing pool was prepared by combining DNA at 2 nM with 12 μl of resuspension buffer containing Tween and 4 μl of diluted PhiX control, then loaded into a P4 flow cell cartridge. Sequencing was performed on an Illumina NextSeq 2000, and run metrics were reviewed on Illumina BaseSpace before analysis. We applied a first prevalence and abundance filter after sequencing to reduce technical noise, retaining species present in more than 5% of samples and with a relative abundance above 0.01%.

Species relative abundances were obtained with MetaPhlAn (55) against the vJan21 CHOCOPhlAn species-level genome bin (SGB) database (mpa_vJan21_CHOCOPhlAnSGB_202103) (56), which assigns reads by clade-specific marker genes and resolves SGBs that include as-yet uncharacterized species; species were retained when present in more than 28 samples and exceeding 0.01% of the cohort’s total reads, resulting in 274 species for the species-level estimation.

### Covariate selection and adjustment

The GAINS clinical record contains over one hundred variables, most of which are medication and comorbidity indicators too rare to adjust for. We drew one adjustment set from that record and used it unchanged in the GAINS species screen, the community ensemble, and the within-community context fits. The ten clinical covariates are an age category, sex, antibiotic use within the past six months, selective serotonin reuptake inhibitor (SSRI) use, proton-pump inhibitor use, statin use, polypharmacy at five or more medications, the Clinical Frailty Scale (57), diabetes, and hypertension, plus visit day as a continuous eleventh term. Three differed between the groups: frailty, SSRI use, and polypharmacy (**Extended Data Table 1**). We excluded cognitive scores, the Clinical Dementia Rating, other dementia coding, and medications prescribed only for Alzheimer’s disease, since each marks the diagnosis rather than confounding it. The unit of analysis is the sample. Each of the 274 metagenomes was treated as one observation; 47 were obtained from 18 participants with AD. No random participant term is fitted. Every covariate except visit day is constant within a participant, so the 274 samples provide more observations than the 119 participants but no more covariate information, and adjustment operates between participants throughout.

### Diversity and composition

Alpha diversity is the Shannon index computed on the species table with vegan (58), compared between groups in a linear mixed model carrying the same ten clinical covariates and visit day with a random intercept for participant, and summarized by the AD coefficient with a Wald 95% confidence interval. Beta diversity is Bray-Curtis distance, tested by permutational multivariate analysis of variance (PERMANOVA) with vegan’s adonis2 over 999 free permutations. The PERMANOVA uses one sample per participant (each participant’s first visit; n = 119), since permuting within a participant cannot shuffle an AD label that is constant within participants. Terms enter sequentially with AD last, so the variance attributed to AD is what remains after the covariates. Sex accounts for 2.3% of the variance in species composition (*P* = 0.001) and AD for 1.0% (*P* = 0.226).

### Differential abundance analysis

The species screen is a single, deterministic LinDA (23) fit over the gut species composition table. The whole profile is estimated jointly in a single call under the shared adjustment set, and because LinDA is fit by least squares with a bias correction for compositionality, the result requires no seed and reproduces exactly. LinDA tests 274 species and returns, for each, an AD log_2_ fold change, its standard error, and a q value from LinDA’s internal Benjamini-Hochberg correction (59), all sorted by q. The primary fit pools all visits, treating visit day as a fixed covariate and excluding a random participant effect, yielding population-averaged estimates. LinDA’s mixed-effect form was not fitted, so the false-discovery control its authors report for correlated samples (23) does not apply, and the screen is a feature-selection step consistent with the sample-level design. We then use a baseline-only fit restricted to each participant’s first visit (visit day 0) as a conservative sensitivity check. For feature selection, we take species passing at q < 0.05 as candidate signals and characterize them in the estimation register by a Cliff’s delta (60) and a carriage prevalence difference, each with participant-level bootstrap intervals. The community ensemble and context dependence that follow are the same LinDA model at finer resolutions. The community ensemble fits it across 1,000 runs at the community level, and the context model fits it within each community across those runs.

### Community modeling across 1,000 random runs

The community ensemble runs in two stages. The first stage is signal generation. We fit latent Dirichlet allocation to the species count matrix by variational expectation-maximization in the topicmodels package (61) using 1,000 independent random seeds, choosing each run’s community count K with the ldatuning package (62) by the CaoJuan2009 (63) and Arun2010 (64) metrics, min-max normalized and averaged, over K from 3 to 100 in steps of 3, yielding K values between 21 and 96 (median 48). Each fit yields K communities and a per-sample membership (gamma) matrix; for each run, we run the same LinDA contrast once on that gamma. Then, across all K communities, we compute an AD log_2_ fold change for each community. As in the species-level analysis, these per-run fits serve as features for feature extraction.

### Matching and collapsing communities across runs

The second stage consolidates across runs. The 48,351 per-run communities (**Fig. 2a,b**) are aligned via nearest-match clustering based on the Jensen-Shannon divergence (65) between their species compositions, with a linear-sum (Hungarian (25)) assignment step used only to build a consensus reference. The clustering retains every community, allowing within-run co-members. The alignment creates a new community when a topic’s Jensen-Shannon divergence from every existing candidate exceeds 0.35, yielding 118 candidate communities. The alignment is label-free. It uses species composition alone, and AD information enters only afterward.

### Community-level differential abundance

A candidate community is considered robust when its AD direction is consistent across runs. Each run casts one up-or-down vote, determined by the sign of its mean fold change. The counts are tested with a chi-square test against a pooled base-rate null, with a Benjamini-Hochberg adjustment and a 70% consistency floor. Finally, the community must recur in more than 500 of the 1,000 runs. Twenty-two are robust, with 11 lower and 11 higher in AD (**Fig. 2c-f** and **Fig. 3**). This test is a selection rule over model runs that all re-fit the same 274 samples. It certifies that a community recurs with a single direction across runs; it is not an inference about the AD association across participants. Any per-species fold change shown for a community comes from the species screen.

### Compositional analyses and reference frames

Pairwise co-variation was assessed using Spearman correlation across all 37,401 unordered pairs of the 274 species, computed from whole-community centered log-ratios of one first-visit sample per participant, with a Benjamini-Hochberg adjustment across all pairs. 12,468 pairs passed at an adjusted probability of 0.05, and their median absolute correlation was 0.34. We characterized the ecology of the candidate signals using a compact set of effect-size analyses. For each candidate species, we estimated an AD-versus-control effect size as Cohen’s *d* (66) on CLR abundance, at the sample level with an analytic 95% interval from the Hedges-Olkin standard error (67) (**Fig. 3a**, **Table 1**) and, as the participant-level check, on the mean CLR per participant, calling a species consistent per person when that interval also excludes zero. Co-occurrence structure was estimated with SPIEC-EASI (68) by neighborhood selection with StARS model selection over 30 regularization values (minimum ratio 0.01, 50 subsamples), which returns 695 conditional relationships among the 260 species that hold at least one. Co-occurrence was reported descriptively as co-abundance rather than as an AD interaction, because the observed network difference did not exceed a participant-level edge-rewiring permutation null. The *P. vulgatus* to *B. fragilis* balance is the difference of their centered log-ratio abundances, one value per first-visit participant, which is algebraically the log-ratio of the two species. The whole-community frame takes the centered log-ratio over the full species profile, with zeros set to half the smallest positive value in that sample; the within-family frame takes it over the 14 prevalent *Bacteroidaceae*. Groups are compared by Cohen’s *d* with a pooled standard deviation and a 95% percentile interval over 2,000 bootstrap resamples of each group. The accompanying probability is a label-free permutation test over 5,000 label shuffles on the area under the curve, scored as the larger of the area and its complement, so it asks whether the balance separates the groups in either direction and cannot support a direction. We also summarized the within-*Bacteroidaceae* relative share and member-species carriage (presence or absence prevalence), each with bootstrap intervals. Lastly, we estimated the co-exclusion of *P. vulgatus* and its closest relative, *P. dorei*, as the Pearson correlation of their CLR abundances, benchmarked against the within-family compositional-closure baseline.

### Indicator species analysis

A classical indicator species analysis was run with indicspecies over all 274 species on the 119 first-visit participants, using the group-size corrected indicator value and the point-biserial correlation (28). Each was run over 9,999 permutations with every species assigned to a single group. The analysis uses neither the community model nor ADAM, so it serves as a convergence check rather than as a biomarker screen.

### Ordination and embedding

The community ensemble is displayed in two dimensions by t-SNE with scikit-learn (69). Each of the 48,351 per-run communities is represented by its composition over the union of the leading 50 species from each run, row-normalized and square-root transformed, then reduced to 50 principal components and embedded with perplexity 40, principal-component initialization, and 1,000 iterations. **Extended Data Fig. 4a** reuses these coordinates rather than re-embedding. **Fig. 3b** is a principal component analysis of the 50 by 22 matrix of species membership weights in the recurring communities, row-normalized, square-root transformed, and mean-centered, with the first two components explaining 17.0% and 11.4% of the variance. Both displays are descriptive, and in the t-SNE, only local neighborhood structure is interpretable.

### External validation in AlzBiom

We tested the candidate signals specified in advance by the GAINS discovery in the independent AlzBiom cohort (ENA PRJEB47976) for AD microbiome ecology (52). AlzBiom comprises 175 stool shotgun metagenomes, one per participant, 100 cognitively healthy controls and 75 participants with probable AD dementia by NIA-AA clinical criteria and cerebrospinal fluid (CSF) Aβ42 confirmation. We re-fetched the raw reads and reprofiled them with MetaPhlAn 3.1.0 (55, 56) against the same vJan21 SGB database used above, so that the two cohorts are comparable at the species level, and included covariates for age, sex, and body mass index (BMI). We performed external validation using effect estimates with 95% confidence intervals within the AD and control arms in AlzBiom. The reported correlations are unadjusted, so both cohorts are compared on the same quantity; the adjusted partial correlations agree to two decimals, –0.57 in GAINS and –0.43 in AlzBiom. The pre-registered gate for the balance was a label-free permutation test on the area under the curve, scored as the larger of the area and its complement, so it asks whether the balance separates the groups in either direction (*P* = 0.038) and cannot support a direction; a permutation on Cohen’s *d* in the GAINS direction gives *P* = 0.121, so the Results rest on the estimate and its interval. We used these external data to check whether the GAINS pattern is AD-associated or reflects general gut ecology.

### The ADAM framework and its literature analytics layer

The ADAM framework (22) couples a vector database (VectorDB), a large language model (LLM), and computational units through retrieval-augmented generation; for this study, we extended it with a species knowledge graph (KG) and managed the data flow among components with a knowledge graph-based retrieval-augmented generation (GraphRAG) approach (70). The VectorDB contains AD-relevant publications from 1983 to 2026, totaling 174,230 publications from 6,321 journals, with most from the past 5 years. The KG contains 11,427 nodes and 19,218 edges spanning all 1,824 GAINS gut species from domain to strain, with each resolved against NCBI Taxonomy (71) as the primary reference and GTDB (72) as a backup for MetaPhlAn genome bins that NCBI cannot name. Each species node carries the species’ taxonomy, name synonyms, and GAINS effect direction (274 species screened, 23 flagged by the species screen). The base LLM used with ADAM is gpt-oss:120b (73), paired with MedEmbed-large-v0.1 (74) to convert publications into VectorDB knowledge (**Extended Data Fig. 7**), and vice versa. Then the computational units, such as topic modeling, LinDA differential abundance analysis, and chi-square tests, provide fundamental bioinformatics support for trustworthy data analytics of the GAINS cohort within the ADAM framework.

For signal identification in the GAINS cohort, we triangulated each candidate signal against the published literature. We mapped each species to its node in a species KG using NCBI taxonomy and incorporated new knowledge from this study. The KG was then linked to the literature from 1983 to 2026 AD via semantic context similarity. The ADAM framework automatically tracks and maps species’ current and historical names to ensure consistency.

### Statistics and reporting

As we define the analysis pipeline for a more exploratory association study, we report estimation statistics throughout, including effect directions, magnitudes, and 95% confidence intervals. The purpose of this design is to generate falsifiable or testable hypotheses for further verification, either by external data or through lab experiments. No statistical method was used to predetermine sample size, which was fixed by the GAINS cohort. One stool sample returned no reads, so we analyzed 274 of the 275 samples. The study is observational, so no randomization was performed, and investigators were not blinded to group during collection or analysis. Sex is recorded in the clinical record as a binary variable and is included as a covariate in every adjusted model; 37 of 101 control participants and 7 of 18 participants with AD are male. Center values are means or fitted model coefficients, as stated in each legend, and intervals are 95% confidence intervals. Bootstrap intervals use 1,000 resamples for correlations and carriage and 2,000 for group means and Cohen’s *d*. Permutation nulls use 999 shuffles for PERMANOVA, 9,999 for the indicator species analysis, and 5,000 for the balance.

### Reproducibility, software, and hardware

We ran analyses on the UMass Chan SCI cluster across CPU (AMD EPYC 7702, Intel Xeon Gold 6230) and GPU (NVIDIA H200, A100, RTX Pro 6000, or V100, as available) nodes. Each of the 1,000 runs selected its own community count K via LDA Tuning rather than using a fixed K, and every random draw was set deterministically by our random seed generator, so the full ensemble reproduces exactly. The Alzheimer’s disease knowledge graph was developed and managed by Obsidian. Analyses used R with LinDA 0.2.0, topicmodels 0.2-16, ldatuning 1.0.2, and the SpiecEasi, vegan, and indicspecies packages, with Python and scikit-learn for the ordinations. MetaPhlAn profiling used the mpa_vJan21_CHOCOPhlAnSGB_202103 database.

### Use of artificial intelligence

The authors disclose the use of artificial intelligence tools in this work. The analysis framework and data visualization code were developed with assistance from Claude Code (Anthropic). The authors conducted literature retrieval and semantic analysis using their in-house ADAM framework, which includes a designated Alzheimer’s disease knowledge graph (**Extended Data Fig. 6**) and an Alzheimer’s disease vector database of 174k publications, built on the gpt-oss:20b and gpt-oss:120b open-weight models within our local repository. These tools also assisted with editing the manuscript, including formatting, reference and citation management, grammar correction, and proofreading. No artificial intelligence tool generated or altered the study’s underlying data, metagenomic profiles, or statistical results, and none is listed as an author or qualifies as an author. The authors reviewed and verified all artificial intelligence-generated output and take full responsibility for the content and conclusions of this manuscript.

## DATA AVAILABILITY

GAINS shotgun metagenomes and de-identified clinical metadata are held by the study group and are available from the corresponding author under a data-use agreement consistent with the governing institutional review board (IRB) protocol (H00021745). AlzBiom metagenomes are publicly available in the European Nucleotide Archive under accession PRJEB47976.

## CODE AVAILABILITY

All analysis code, including ADAM and MT-LDA implementation, is archived at Zenodo (https://doi.org/10.5281/zenodo.22682330), with per-figure source-data tables and per-stage documentation.

## ACKNOWLEDGMENT

We thank the participants of the GAINS cohort and their families for their contributions, which made this work possible. We thank the members of the Haran Laboratory at the University of Massachusetts Chan Medical School for coordinating clinical and microbiome studies and collecting samples. We also acknowledge the Bucci Laboratory for technical guidance and feedback that shaped the compositional and community-level analyses. All analyses were conducted in-house at the University of Massachusetts Chan Medical School.

## FUNDING

**John P. Haran** and **Vanni Bucci** disclose support for the research of this work from the National Institute on Aging [5R01AG075283-04, 2R01AG067483-03].

## AUTHOR CONTRIBUTION

**Z.H., B.A.M.** and **J.P.H.** conceptualized the study. **Z.H., V.B.** and **J.P.H.** developed the methodology. **Z.H., D.V.W.** and **V.B.** performed the formal analysis. **Z.H.** developed the software and prepared the visualizations. **P.M.M., D.C.F.** and **J.P.H.** curated the data. **P.M.M.** conducted clinical data investigation and management. **D.C.F., B.A.M., D.V.W., V.B.** and **J.P.H.** contributed to the investigation. **V.B.** and **J.P.H.** acquired funding. **B.A.M., V.B.** and **J.P.H.** supervised the study. **Z.H.** and **J.P.H.** wrote the original draft**. Z.H., D.C.F., B.A.M., D.V.W., V.B.** and **J.P.H.** reviewed and edited the manuscript.

## COMPETING INTERESTS

The authors declare no competing interests.

## EXTENDED DATA

**Extended Data Fig. 1.**
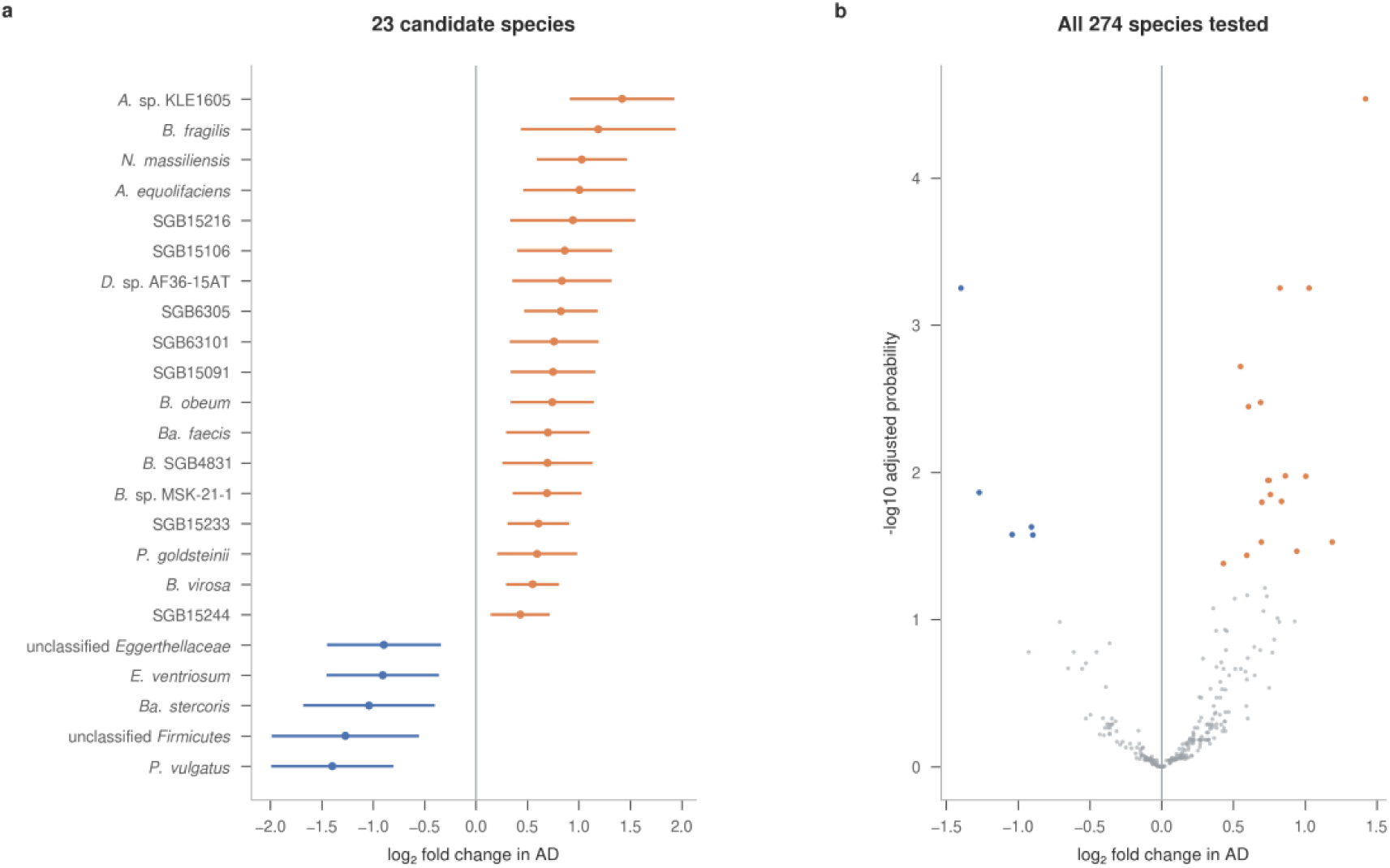
Species-level differential abundance in Alzheimer’s disease. **a**, Covariate-adjusted log_2_ fold change with its 95% confidence interval for the 23 candidate species, colored by direction (blue, lower in AD; orange, higher in AD). The largest enrichment is *Akkermansia* sp. KLE1605 (+1.42) and the sharpest depletion is *Phocaeicola vulgatus* (–1.40), the most abundant species in the cohort. **b,** Volcano plot of log_2_ fold change against –log_10_ Benjamini-Hochberg adjusted probability for all 274 species tested, with 23 candidates colored by direction and the remainder in gray. Candidates are the 23 species below the screening threshold of a Benjamini-Hochberg adjusted probability of 0.05. n = 274 samples from 119 participants, 18 with AD. Center values are LinDA-fitted log2 fold changes adjusted for the ten clinical covariates and visit day listed in Methods; intervals are 95% confidence intervals.

**Extended Data Fig. 2.**
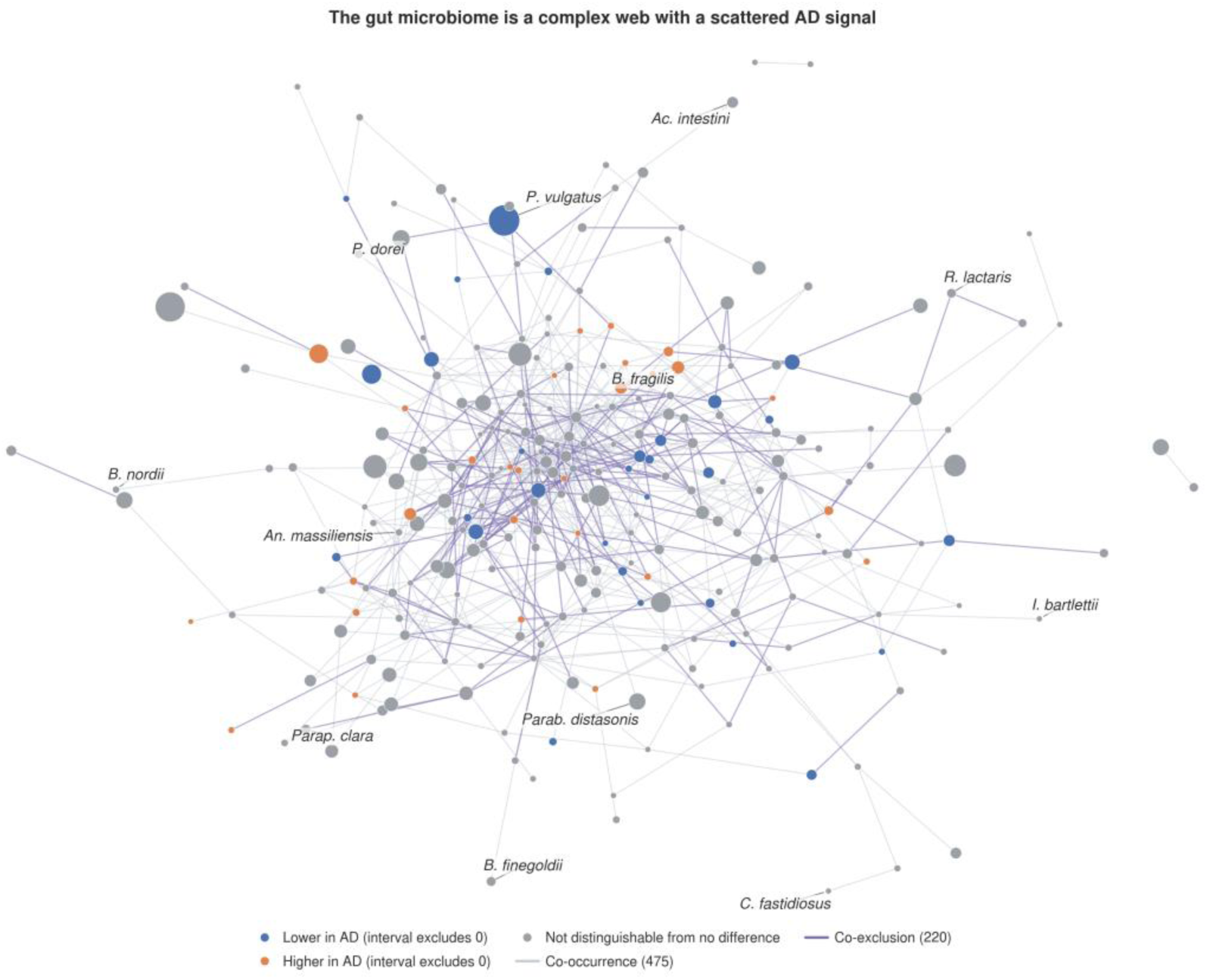
The species that differ in Alzheimer’s disease are scattered across the gut co-abundance network. Each node is one of the 260 gut species holding at least one abundance relationship that survives conditioning on every other species, placed by force-directed layout and sized by mean relative abundance. A node carries a direction color only where the 95% confidence interval on its sample-level AD effect excludes zero, a wider criterion than the species screen behind **Extended Data** Fig. 1 (blue, lower in AD, 29 species; orange, higher in AD, 28 species), and the remaining 203 species are gray. Each of the 695 lines is one such relationship, drawn in pale gray for the 475 co-occurrences and in purple for the 220 co-exclusions. A few species are named for orientation, including *Phocaeicola vulgatus*, *Phocaeicola dorei,* and *Bacteroides fragilis*. We estimated the network across all 274 samples. Species that differ in AD occupy no particular region of the network and hold no special position within it, so the disease signal is distributed across the community rather than concentrated in one part. Node degree is descriptive only, since hub structure can arise under the null in an all-pairs screen and is not evidence that a species organizes the community. n = 274 samples from 119 participants, 18 with AD.

**Extended Data Fig. 3.**
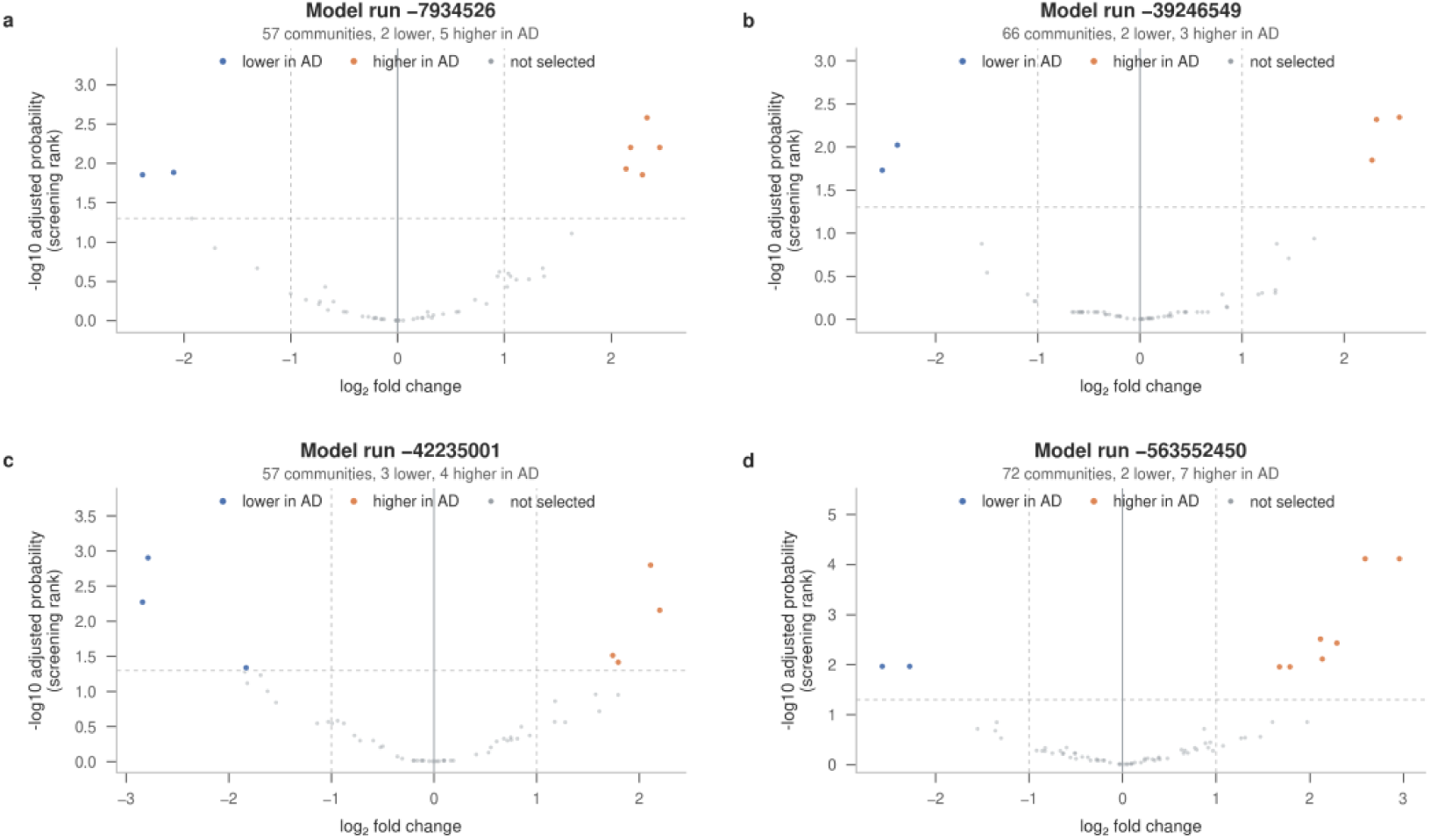
Community structure varies from one model run to the next. **a** to **d,** Community-level volcanoes from four individual model runs, each plotting the community-level log_2_ fold change against –log_10_ Benjamini-Hochberg adjusted probability. The four runs resolved 57, 66, 57, and 72 communities. Communities flagged by the per-run screen are colored by direction (blue, lower in AD; orange, higher in AD), and the remainder are gray; the count in each panel gives the number of flagged communities in each direction. Latent Dirichlet allocation is fit from a random starting point, so different runs recover similar but not identical communities. n = 274 samples from 119 participants in every run. The alignment in **Extended Data** Fig. 4 is what resolves this run-to-run variability.

**Extended Data Fig. 4.**
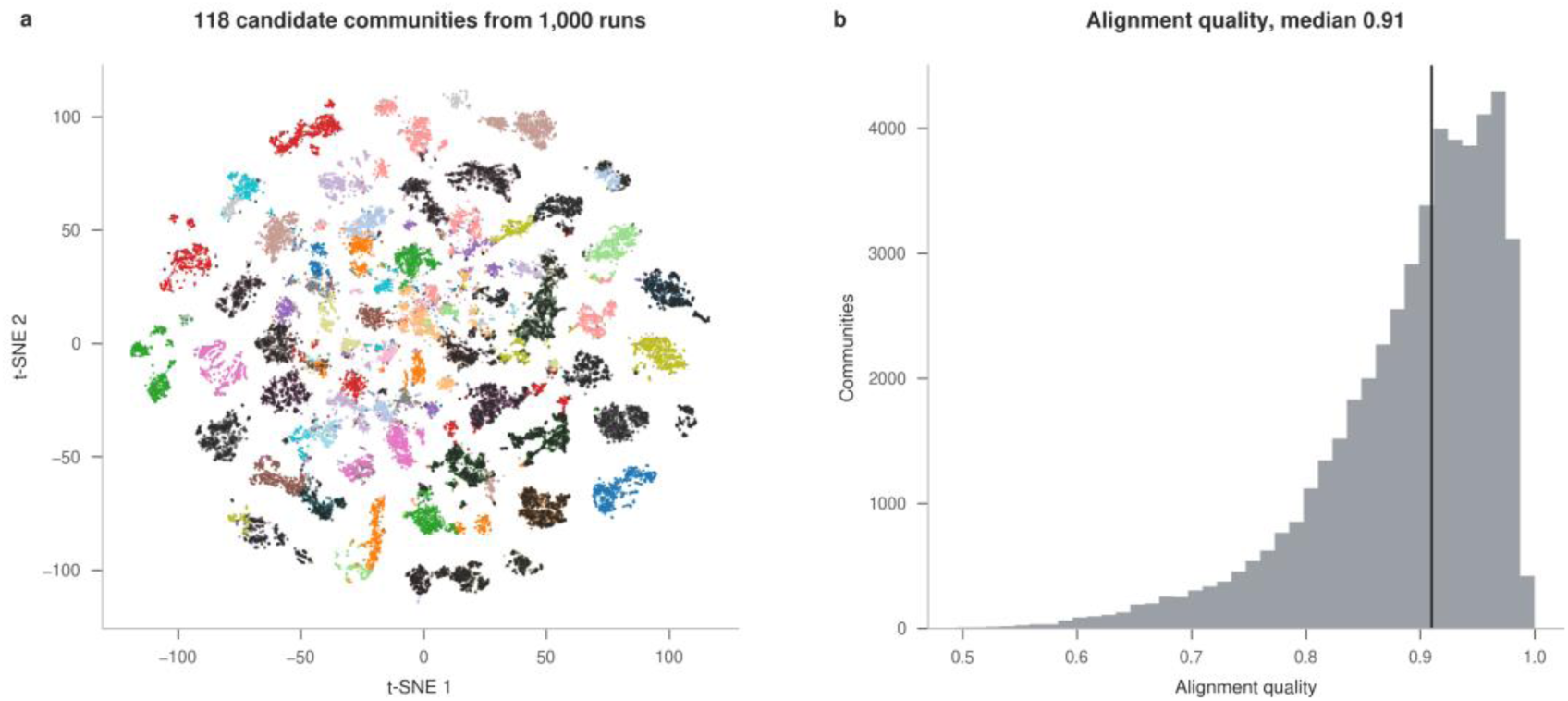
Alignment of the community ensemble into 118 candidates. **a**, All 48,351 per-run communities from the 1,000 model runs, placed on the compositional-similarity embedding and colored by the candidate community each was assigned to. The palette holds 20 colors and cycles across the 118 candidates, so a repeated color does not mark a shared community. A darker layer over 17,727 points (37%) marks the per-run communities belonging to the 22 that recur. Both t-SNE axes are unitless and only local neighborhood composition is interpretable; inter-island distances and island sizes are not. **b,** Distribution of alignment quality, one minus the Jensen-Shannon divergence from the assigned candidate, scaled by log 2, for 48,351 per-run communities, with the vertical line at the median of 0.91. Both the embedding and the alignment are derived from species composition, so **panel a** serves as a consistency check rather than as independent validation.

**Extended Data Fig. 5.**
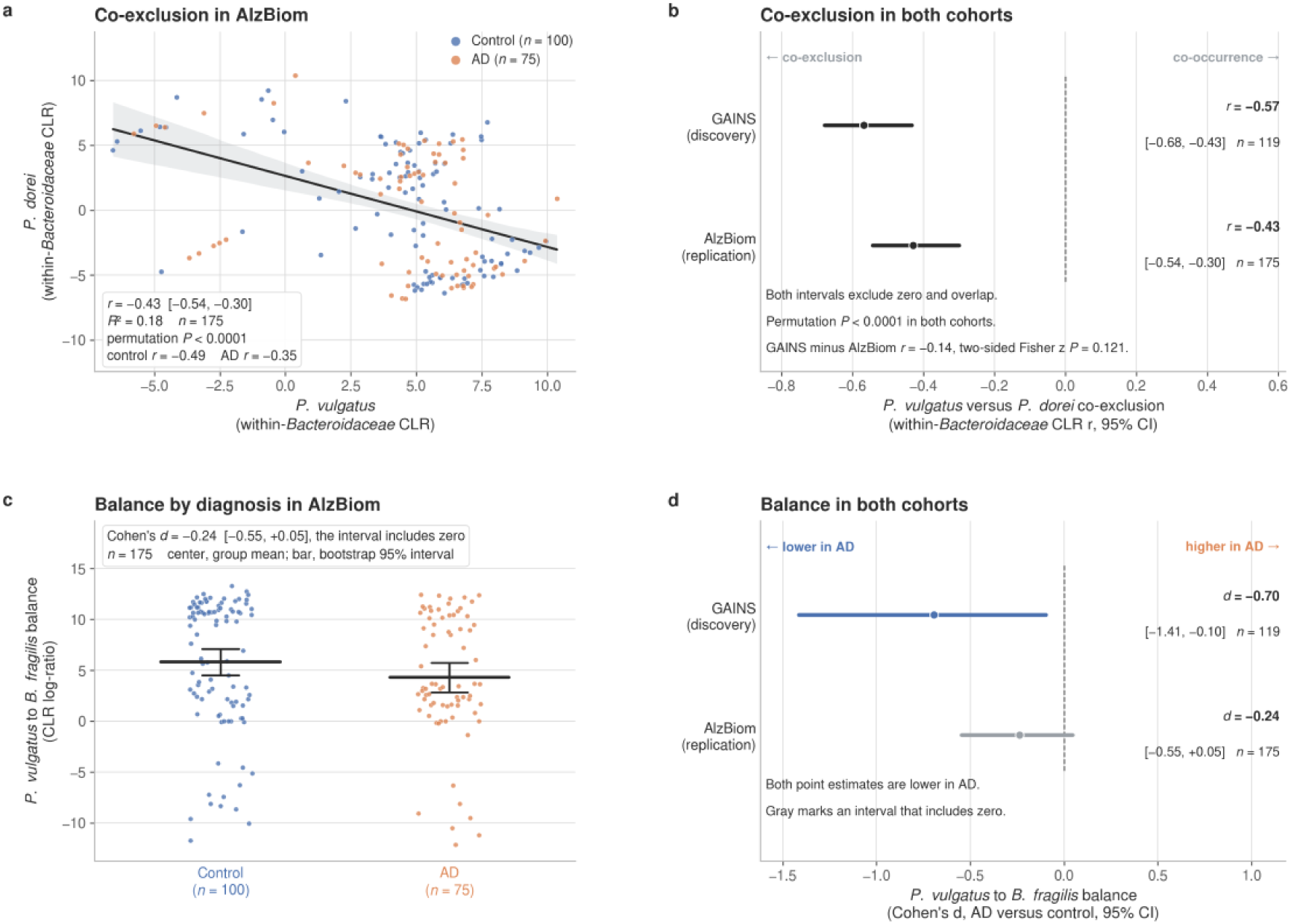
The co-exclusion reproduces in an independent cohort, and the balance repeats its direction only. **a**, Within-Bacteroidaceae centered log-ratio abundance of *P. dorei* against *P. vulgatus* in the independent AlzBiom cohort, one point per participant, colored by diagnosis, control (n = 100) and AD (n = 75). The line is the least-squares fit, and the band is its 95% bootstrap interval over 1,000 resamples. Pearson r = –0.43, 95% CI [-0.54, –0.30] by Fisher z, label-free permutation P < 0.0001; control r = –0.49, AD r = –0.35. **b,** The same correlation in both cohorts, GAINS r = –0.57, 95% CI [-0.68, –0.43], n = 119, and AlzBiom r = –0.43, n = 175, with permutation P < 0.0001 in each. The intervals exclude zero and overlap, and the difference between them is –0.14; two-sided Fisher z P = 0.121. **c,** The *P. vulgatus* to *B. fragilis* balance by diagnosis in AlzBiom. Horizontal bars are group means and vertical bars are their bootstrap 95% confidence intervals over 2,000 resamples. Cohen’s d = –0.24, 95% CI [-0.55, +0.05], which includes zero. **d,** The balance in both cohorts: GAINS d = –0.70, 95% CI [-1.41, –0.10], and AlzBiom in gray to mark an interval that includes zero. Both point estimates are lower in AD. The label-free permutation gate is computed on the area under the curve and is direction-agnostic.

**Extended Data Fig. 6.**
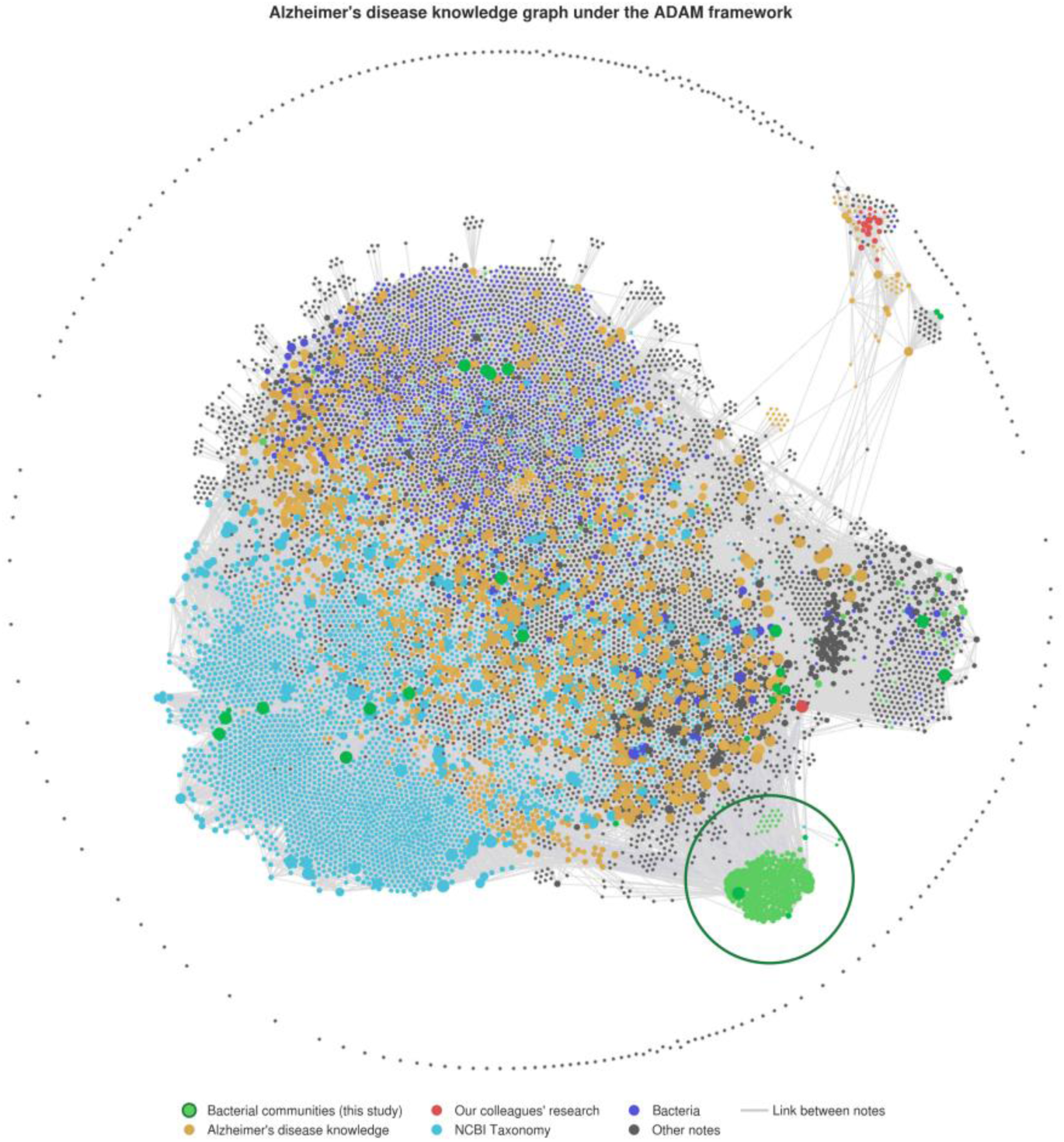
Alzheimer’s disease knowledge graph under the ADAM framework. The knowledge graph organizes all 1,824 gut species profiled in GAINS against the NCBI Taxonomy hierarchy, with the Genome Taxonomy Database naming the genome bins that NCBI cannot. Each species node includes its taxonomy, name synonyms, and its position in the GAINS species screen, and links to the Alzheimer’s disease literature the framework holds in its vector database, 174,230 publications from 6,321 journals published between 1983 and 2026 and retrieved from PubMed Central. The graph and corpus are stored in a local repository, so retrieval runs without an external service. The graph enables ADAM to place a species alongside what has already been reported about it, and it is the source of the key references in **Table 1**. It carries no effect estimate of its own, and nothing in it was used to compute the differences reported in the results. The knowledge graph is maintained and managed using Obsidian. Node color marks the kind of note: **green** for the bacterial communities identified in this study, with the **dark green** circle outlining their cluster; **yellow** for Alzheimer’s disease knowledge; **red** for our colleagues’ research; **light blue** for NCBI Taxonomy; **deep blue** for bacteria; and **gray** for every other note; **gray lines** are links between notes.

**Extended Data Fig. 7.**
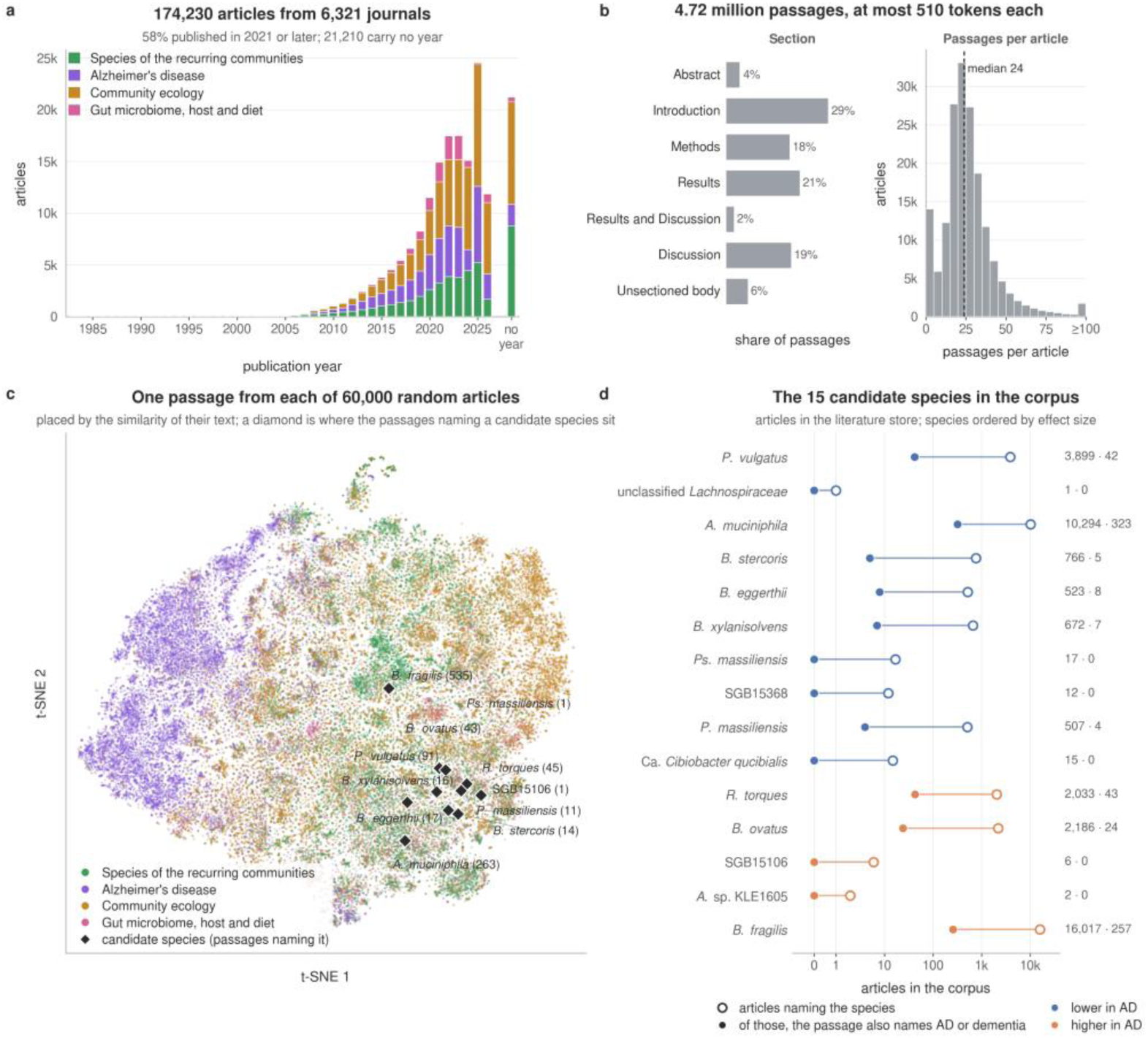
The literature layer of the ADAM framework. The framework stores the published literature as text passages with numerical embeddings; the knowledge graph in **Extended Data** Fig. 6 draws its references from it. **a,** The 174,230 articles by publication year, colored by the search that retrieved each from PubMed Central (Alzheimer’s disease, the species of the 22 recurring communities, community ecology, or the gut microbiome, host and diet). The 21,210 articles without a year form their own bar; 58% of all articles date from 2021 or later. **b,** Each article is split into passages of at most 510 tokens, for a total of 4.72 million. The bars give the share of passages in each section of an article, and the histogram shows passages per article, with a median of 24. **c,** One random body passage from each of 60,000 random articles, placed by the similarity of its 1,024-dimensional MedEmbed embedding (principal components reduced to 50 dimensions, then t-SNE) and colored as in a. Each black diamond marks the median position of the sampled passages that name one of the 15 community species that differ in AD, with their count in parentheses; a species named by no sampled passage has no diamond. **d,** Coverage of the same 15 species within the store, ordered by effect size and colored by direction in AD. The open circle counts articles in which a passage names the species, under its current name or any earlier name; the filled circle counts articles in which such a passage also names Alzheimer’ disease or dementia. Six of the 15, four of them unnamed genome bins or strain-level clades, appear in fewer than 20 articles each and never alongside AD. Counts of text, not estimates of effect.

**Extended Data Fig. 8.**
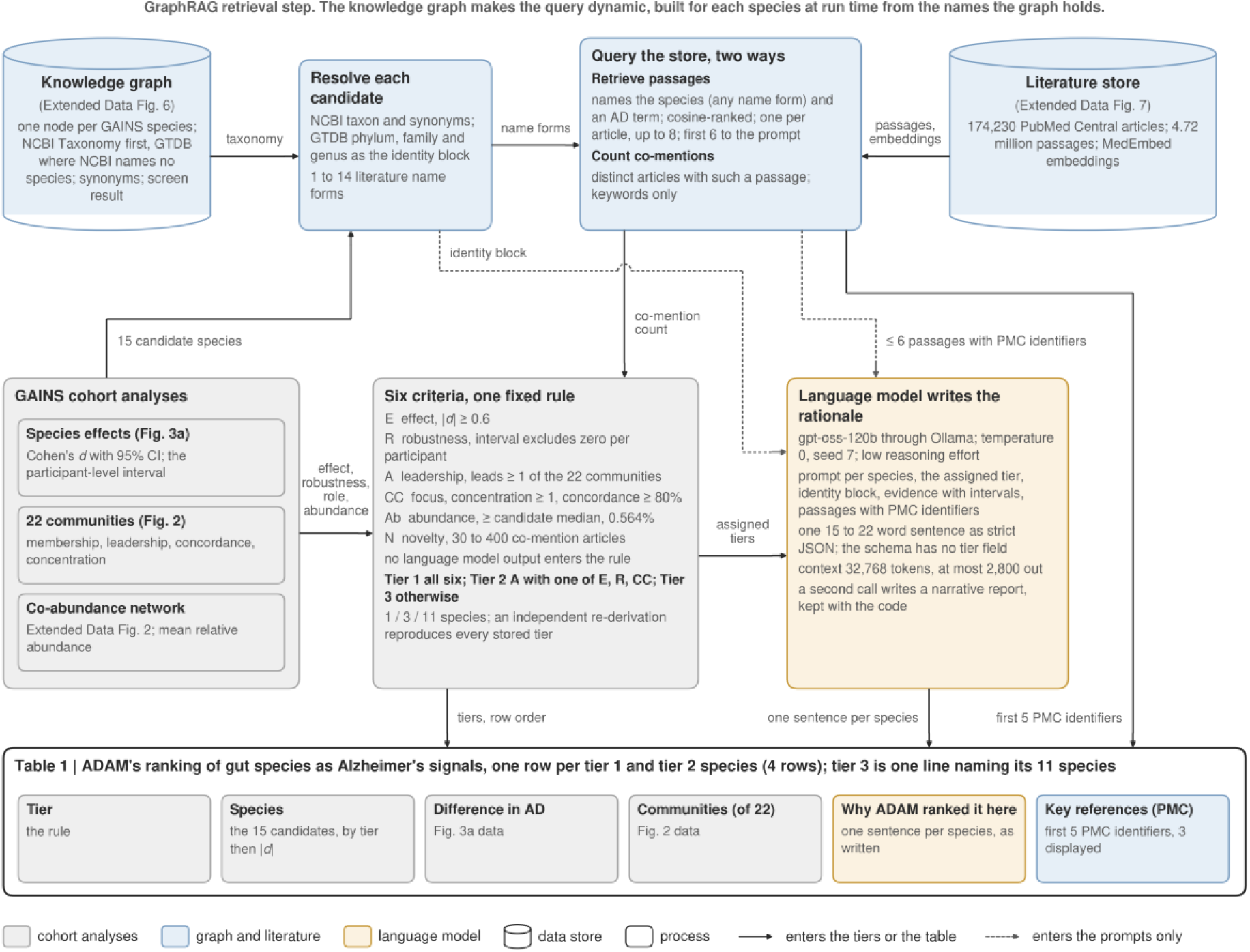
Data flow of the ADAM tier assignment behind Table 1. The ADAM framework ranks species by data flow. Cylinders represent two stores, the knowledge graph (**Extended Data** Fig. 6) and the literature store (**Extended Data** Fig. 7). Rounded boxes indicate processes, and each arrow is labeled with the data it transmits. Fill marks the component, gray the cohort analyses, blue the knowledge graph and the literature, and gold the language model. Solid arrows carry data into the tier assignment or the table; dashed arrows carry data into the language model’s prompts only. The top row is the retrieval step of GraphRAG, retrieval-augmented generation guided by a knowledge graph, and the language model is its generation step. The knowledge graph is the key component, since it makes the query dynamic, built for each species at run time rather than from a fixed search string. Each of the 15 candidate species (Fig. 3a) is resolved in the graph to its literature name forms, from NCBI Taxonomy, where NCBI names the species, and to an identity block of its GTDB phylum, family, and genus. The literature store is then queried under those name forms, so text written under an earlier name, such as *Bacteroides vulgatus* for *Phocaeicola vulgatus*, is retrieved. One query retrieves passages that name the species and an Alzheimer’s-context term, ranked by embedding similarity; the other counts the articles holding such a passage. The bottom row is the ranking. Six criteria with fixed thresholds and one rule determine the tiers, yielding one species in tier 1, three in tier 2, and 11 in tier 3. An independent re-derivation of the rule reproduces all stored tiers. The language model receives the assigned tiers as input, writes one sentence per species, and its output schema has no tier field, so no dashed arrow points at a tier.

**Extended Data Table 1.** GAINS cohort characteristics and covariate balance. Values are mean ± s.d. for continuous variables and n (%) for categorical variables, in 119 participants contributing 274 samples, 18 with AD. *P* values are from two-sided Wilcoxon rank-sum tests for continuous variables and Fisher’s exact tests for categorical variables, without adjustment for multiple comparisons; they describe covariate balance and are not tests of the study hypothesis. Frailty, selective serotonin reuptake inhibitor use, and polypharmacy differ between groups; all ten characteristics, with visit day, are carried as covariates in every adjusted model.

| Group | Characteristic | Control (n = 101) | AD (n = 18) | P value |
| --- | --- | --- | --- | --- |
| Demographics | Age category (1–3) | 1.19 ± 0.76 | 1.56 ± 0.62 | 0.051 |
|  | Male sex | 37 (37) | 7 (39) | 1.000 |
| Geriatric / host state | Clinical Frailty Scale (1–9) | 2.13 ± 0.97 | 3.39 ± 1.46 | < 0.001 |
| Medications | SSRIs | 14 (14) | 8 (44) | 0.005 |
|  | Polypharmacy (≥5) | 27 (27) | 12 (67) | 0.002 |
|  | Antibiotics (≤6 mo) | 19 (19) | 2 (11) | 0.521 |
|  | Proton-pump inhibitors | 18 (18) | 5 (28) | 0.338 |
|  | Statins | 45 (45) | 10 (56) | 0.447 |
| Comorbidities | Hypertension | 45 (45) | 12 (67) | 0.123 |
|  | Diabetes | 10 (10) | 1 (6) | 1.000 |

Values are mean ± SD (continuous) or n (%) (categorical). P values compare control against AD by two-sided Wilcoxon rank-sum (Mann-Whitney U) tests (continuous) or Fisher's exact test (categorical). One record per participant (first visit). 119 participants (101 control, 18 AD) contributed 274 fecal metagenomes; 73 gave ≥2 samples. The one empty AD sample (JPH050S90) is excluded, and differential-abundance models are fit on all 274 samples with these covariates adjusted.

**Extended Data Table 2.** Both *Bacteroidaceae* relationships under two compositional reference frames. We recompute each relationship from raw abundances in each cohort under the within-*Bacteroidaceae* CLR over the 14 prevalent family members and under the whole-community CLR over the full species profile. Raw relative abundance is omitted because untransformed proportions carry the constant-sum constraint and are not a valid compositional treatment. The *P. vulgatus* / *P. dorei* co-exclusion is the Pearson correlation of the two congeners’ abundances with a Fisher 95% confidence interval. The *P. vulgatus* / *B. fragilis* balance is Cohen’s *d* of their compositional log-ratio between AD and control participants with a bootstrap 95% confidence interval. GAINS is the discovery cohort (n = 119 first-visit participants, 18 with AD); AlzBiom is an independent Alzheimer’s case-control cohort (n = 175). The co-exclusion is negative under both frames in both cohorts. The balance is lower in AD under both frames in both cohorts, and in GAINS, its intervals exclude zero, whereas in AlzBiom they include zero.

| Relationship | Cohort | Within-*Bacteroidaceae* CLR | Whole-community CLR |
| --- | --- | --- | --- |
| *P. vulgatus* - *P. dorei* co-exclusion (Pearson r) | GAINS | -0.57 (-0.68, -0.43) | -0.36 (-0.51, -0.19) |
|  | AlzBiom | -0.43 (-0.54, -0.30) | -0.21 (-0.35, -0.07) |
| *P. vulgatus* / *B. fragilis* balance (Cohen's d) | GAINS | -0.67 (-1.41, -0.05) | -0.70 (-1.41, -0.10) |
|  | AlzBiom | -0.23 (-0.53, +0.05) | -0.24 (-0.55, +0.05) |

## Notes

### Competing Interest Statement

The authors have declared no competing interest.

### Summary of Updates

We revised the manuscript in response to the co-authors' feedback and comments, editing the paragraphs and adding new figures.

## References

1. Alzheimer’s A. 2025 Alzheimer’s Disease Facts and Figures. Alzheimer’s & Dementia. 2025;21(5):872–1019.

2. Cattaneo A, Cattane N, Galluzzi S, Provasi S, Lopizzo N, Festari C, et al. Association of brain amyloidosis with pro-inflammatory gut bacterial taxa and peripheral inflammation markers in cognitively impaired elderly. Neurobiol Aging. 2017;49:60–8.

3. Shukla PK, Delotterie DF, Xiao J, Pierre JF, Rao R, McDonald MP, et al. Alterations in the Gut-Microbial-Inflammasome-Brain Axis in a Mouse Model of Alzheimer’s Disease. Cells. 2021;10(4).

4. Wu ML, Yang XQ, Xue L, Duan W, Du JR. Age-related cognitive decline is associated with microbiota-gut-brain axis disorders and neuroinflammation in mice. Behav Brain Res. 2021;402:113125.

5. Xia Y, Xiao Y, Wang ZH, Liu X, Alam AM, Haran JP, et al. Bacteroides Fragilis in the gut microbiomes of Alzheimer’s disease activates microglia and triggers pathogenesis in neuronal C/EBPβ transgenic mice. Nat Commun. 2023;14(1):5471.

6. Bairamian D, Sha S, Rolhion N, Sokol H, Dorothée G, Lemere CA, et al. Microbiota in neuroinflammation and synaptic dysfunction: a focus on Alzheimer’s disease. Mol Neurodegener. 2022;17(1):19.

7. Li B, He Y, Ma J, Huang P, Du J, Cao L, et al. Mild cognitive impairment has similar alterations as Alzheimer’s disease in gut microbiota. Alzheimers Dement. 2019;15(10):1357–66.

8. Ling Z, Zhu M, Yan X, Cheng Y, Shao L, Liu X, et al. Structural and Functional Dysbiosis of Fecal Microbiota in Chinese Patients With Alzheimer’s Disease. Front Cell Dev Biol. 2020;8:634069.

9. Zhang T, Gao G, Kwok L, Sun Z. Gut microbiome-targeted therapies for Alzheimer’s disease. Gut Microbes. 2023;15.

10. Seo D-O, Holtzman DM. Current understanding of the Alzheimer’s disease-associated microbiome and therapeutic strategies. Experimental & Molecular Medicine. 2024;56:86–94.

11. Coyte KZ, Rakoff-Nahoum S. Understanding Competition and Cooperation within the Mammalian Gut Microbiome. Curr Biol. 2019;29(11):R538–r44.

12. Faust K, Sathirapongsasuti JF, Izard J, Segata N, Gevers D, Raes J, et al. Microbial co-occurrence relationships in the human microbiome. PLoS Comput Biol. 2012;8(7):e1002606.

13. Vogt NM, Kerby RL, Dill-McFarland KA, Harding SJ, Merluzzi AP, Johnson SC, et al. Gut microbiome alterations in Alzheimer’s disease. Scientific Reports. 2017;7(1):13537.

14. Blei DM, Ng AY, Jordan MI. Latent dirichlet allocation. J Mach Learn Res. 2003;3(null):993–1022.

15. Jelodar H, Wang Y, Yuan C, Feng X, Jiang X, Li Y, et al. Latent Dirichlet allocation (LDA) and topic modeling: models, applications, a survey. Multimedia Tools and Applications. 2019;78(11):15169–211.

16. Fitzjerrells RL, Ollberding NJ, Mangalam AK. Looking at the full picture, using topic modeling to observe microbiome communities associated with disease. Gut Microbes Rep. 2024;1(1):1–11.

17. Tataru C, Peras M, Rutherford E, Dunlap K, Yin X, Chrisman BS, et al. Topic modeling for multi-omic integration in the human gut microbiome and implications for Autism. Sci Rep. 2023;13(1):11353.

18. Rieger J, Jentsch C, Rahnenführer J. LDAPrototype: a model selection algorithm to improve reliability of latent Dirichlet allocation. PeerJ Comput Sci. 2024;10:e2279.

19. Mantyla MV, Claes M, Farooq U. Measuring LDA topic stability from clusters of replicated runs. Proceedings of the 12th ACM/IEEE International Symposium on Empirical Software Engineering and Measurement; Oulu, Finland: Association for Computing Machinery; 2018. p. Article 49.

20. Giampino A, Ascari R, Migliorati S. A flexible mixed-membership model for community and enterotype detection for microbiome data. Comput Stat Data Anal. 2025;210(C):15.

21. Haran JP, Barrett AM, Lai Y, Odjidja SN, Dutta P, McGrath PM, et al. Executive functioning and processing speed as predictors of global cognitive decline in Alzheimer’s disease. J Alzheimers Dis Rep. 2025;9:25424823251363549.

22. Huang Z, Kaur Sekhon V, Sadeghian R, Vaida ML, Jo C, McCormick BA, et al. ADAM-1: An AI Reasoning and Bioinformatics Model for Alzheimer’s Disease Detection and Microbiome-Clinical Data Integration. IEEE Access. 2025;13:145953–67.

23. Zhou H, He K, Chen J, Zhang X. LinDA: linear models for differential abundance analysis of microbiome compositional data. Genome Biol. 2022;23(1):95.

24. Sankaran K, Holmes SP. Latent variable modeling for the microbiome. Biostatistics. 2019;20(4):599–614.

25. Kuhn HW. The Hungarian method for the assignment problem. Naval research logistics quarterly. 1955;2(1-2):83–97.

26. Clausen U, Vital ST, Lambertus P, Gehler M, Scheve S, Wöhlbrand L, et al. Catabolic Network of the Fermentative Gut Bacterium Phocaeicola vulgatus (Phylum Bacteroidota) from a Physiologic-Proteomic Perspective. Microbial physiology. 2024;34(1):88–107.

27. Gindt ME, Lück R, Deppenmeier U. Genetic optimization of the human gut bacterium Phocaeicola vulgatus for enhanced succinate production. Applied microbiology and biotechnology. 2024;108(1):465.

28. De Cáceres M, Legendre P. Associations between species and groups of sites: indices and statistical inference. Ecology. 2009;90(12):3566–74.

29. Jean S, Wallace Miranda J, Dantas G, Burnham Carey-Ann D. Time for Some Group Therapy: Update on Identification, Antimicrobial Resistance, Taxonomy, and Clinical Significance of the Bacteroides fragilis Group. Journal of Clinical Microbiology. 2022;60(9):e02361–20.

30. Tajkarimi M, Wexler HM. CRISPR-Cas Systems in Bacteroides fragilis, an Important Pathobiont in the Human Gut Microbiome. Frontiers in Microbiology. 2017;Volume 8 – 2017.

31. Rivera-Pinto J, Egozcue JJ, Pawlowsky-Glahn V, Paredes R, Noguera-Julian M, Calle ML. Balances: a New Perspective for Microbiome Analysis. mSystems. 2018;3(4).

32. Cabrera C, Carrión N, Mateo D, Vicens P, Pinzón A, Heredia L, et al. Gut microbiota characterization in ageing, mild cognitive impairment, and Alzheimer’s disease in the context of mediterranean lifestyle in a Spanish population. Alzheimers Res Ther. 2025;17(1):211.

33. De Jaegher S, Pinzauti D, D’Aguanno M, Parkinson E, Schofield J, Strazzeri F, et al. Identifying microbial biomarkers of neurodegeneration: a comparative study in Alzheimer’s and Parkinson’s disease. Front Microbiomes. 2026;5:1831956.

34. Han Y, Quan X, Chuang Y, Liang Q, Li Y, Yuan Z, et al. A multi-omics analysis for the prediction of neurocognitive disorders risk among the elderly in Macao. Clin Transl Med. 2022;12(6):e909.

35. Haran JP, Bhattarai SK, Foley SE, Dutta P, Ward DV, Bucci V, et al. Alzheimer’s Disease Microbiome Is Associated with Dysregulation of the Anti-Inflammatory P-Glycoprotein Pathway. mBio. 2019;10(3).

36. Jahvnavi, Garg P, Srivastava P. Metagenomics study reveals altered composition of Avispirillum, Phocaeicola, Bacteroides, and Faecalibacterium in the gut of patients suffering with Alzheimer’s disease. Ecological Genetics and Genomics. 2025;34:100322.

37. Park S, Wu X. Modulation of the Gut Microbiota in Memory Impairment and Alzheimer’s Disease via the Inhibition of the Parasympathetic Nervous System. Int J Mol Sci. 2022;23(21).

38. Son SJ, Wu X, Roh HW, Cho YH, Hong S, Nam YJ, et al. Distinct gut microbiota profiles and network properties in older Korean individuals with subjective cognitive decline, mild cognitive impairment, and Alzheimer’s disease. Alzheimers Res Ther. 2025;17(1):187.

39. Zhang D, Jian YP, Zhang YN, Li Y, Gu LT, Sun HH, et al. Short-chain fatty acids in diseases. Cell Commun Signal. 2023;21(1):212.

40. Popescu C, Munteanu C, Anghelescu A, Ciobanu V, Spînu A, Andone I, et al. Novelties on Neuroinflammation in Alzheimer’s Disease-Focus on Gut and Oral Microbiota Involvement. Int J Mol Sci. 2024;25(20).

41. Da Silva Morais E, Grimaud GM, Warda A, Stanton C, Ross P. Genome plasticity shapes the ecology and evolution of Phocaeicola dorei and Phocaeicola vulgatus. Sci Rep. 2024;14(1):10109.

42. Yunusbaeva M, Borodina L, Terentyeva D, Bogdanova A, Zakirova A, Bulatov S, et al. Excess fermentation and lactic acidosis as detrimental functions of the gut microbes in treatment-naive TB patients. Front Cell Infect Microbiol. 2024;14:1331521.

43. Yan A, Butcher J, Mack DR, Stintzi A. The colonic mucosal virome in inflammatory bowel disease reveals Crassvirales depletion and disease-specific virome features. Gut Microbes. 2025;17(1):2539450.

44. Wexler Hannah M. Bacteroides: the Good, the Bad, and the Nitty-Gritty. Clinical Microbiology Reviews. 2007;20(4):593–621.

45. Sun F, Zhang Q, Zhao J, Zhang H, Zhai Q, Chen W. A potential species of next-generation probiotics? The dark and light sides of Bacteroides fragilis in health. Food Res Int. 2019;126:108590.

46. Shin JH, Tillotson G, MacKenzie TN, Warren CA, Wexler HM, Goldstein EJC. Bacteroides and related species: The keystone taxa of the human gut microbiota. Anaerobe. 2024;85:102819.

47. Qu D, Sun F, Feng S, Yu L, Tian F, Zhang H, et al. Protective effects of Bacteroides fragilis against lipopolysaccharide-induced systemic inflammation and their potential functional genes. Food Funct. 2022;13(2):1015–25.

48. Valguarnera E, Wardenburg JB. Good Gone Bad: One Toxin Away From Disease for Bacteroides fragilis. J Mol Biol. 2020;432(4):765–85.

49. Wasén C, Beauchamp LC, Vincentini J, Li S, LeServe DS, Gauthier C, et al. Bacteroidota inhibit microglia clearance of amyloid-beta and promote plaque deposition in Alzheimer’s disease mouse models. Nat Commun. 2024;15(1):3872.

50. Lukiw WJ. Bacteroides fragilis Lipopolysaccharide and Inflammatory Signaling in Alzheimer’s Disease. Front Microbiol. 2016;7:1544.

51. Popescu C, Munteanu C, Anghelescu A, Ciobanu V, Spînu A, Andone I, et al. Novelties on Neuroinflammation in Alzheimer’s Disease–Focus on Gut and Oral Microbiota Involvement. International Journal of Molecular Sciences. 2024;25.

52. Laske C, Müller S, Preische O, Ruschil V, Munk M, Honold I, et al. Signature of Alzheimer’s Disease in Intestinal Microbiome: Results From the AlzBiom Study. Frontiers in neuroscience. 2022.

53. Engevik AC, Engevik MA. Exploring the impact of intestinal ion transport on the gut microbiota. Comput Struct Biotechnol J. 2021;19:134–44.

54. Jack Jr CR, Bennett DA, Blennow K, Carrillo MC, Dunn B, Haeberlein SB, et al. NIA-AA Research Framework: Toward a biological definition of Alzheimer’s disease. Alzheimer’s & Dementia. 2018;14(4):535–62.

55. Beghini F, McIver LJ, Blanco-Míguez A, Dubois L, Asnicar F, Maharjan S, et al. Integrating taxonomic, functional, and strain-level profiling of diverse microbial communities with bioBakery 3. Elife. 2021;10.

56. Blanco-Míguez A, Beghini F, Cumbo F, McIver LJ, Thompson KN, Zolfo M, et al. Extending and improving metagenomic taxonomic profiling with uncharacterized species using MetaPhlAn 4. Nat Biotechnol. 2023;41(11):1633–44.

57. Rockwood K, Song X, MacKnight C, Bergman H, Hogan DB, McDowell I, et al. A global clinical measure of fitness and frailty in elderly people. Cmaj. 2005;173(5):489–95.

58. Oksanen J, Simpson GL, Blanchet FG, Kindt R, Legendre P, Minchin PR, et al. vegan: Community Ecology Package. R package version 2.7-3 ed: Comprehensive R Archive Network (CRAN); 2026.

59. Benjamini Y, Hochberg Y. Controlling the False Discovery Rate: A Practical and Powerful Approach to Multiple Testing. Journal of the Royal Statistical Society: Series B (Methodological). 1995;57(1):289–300.

60. Cliff N. Dominance statistics: Ordinal analyses to answer ordinal questions. Psychological bulletin. 1993;114(3):494.

61. Grün B, Hornik K. topicmodels: An R Package for Fitting Topic Models. Journal of Statistical Software. 2011;40(13):1–30.

62. Murzintcev N. ldatuning: Tuning of the Latent Dirichlet Allocation Models Parameters. R package version 1.0.2 ed: Comprehensive R Archive Network (CRAN); 2020.

63. Cao J, Xia T, Li J, Zhang Y, Tang S. A density-based method for adaptive LDA model selection. Neurocomputing. 2009;72(7):1775–81.

64. Arun R, Suresh V, Veni Madhavan CE, Narasimha Murthy MN, editors. On Finding the Natural Number of Topics with Latent Dirichlet Allocation: Some Observations 2010; Berlin, Heidelberg: Springer Berlin Heidelberg.

65. Lin J. Divergence measures based on the Shannon entropy. IEEE Transactions on Information Theory. 1991;37(1):145–51.

66. Cohen J. Statistical power analysis for the behavioral sciences: routledge; 2013.

67. Hedges Larry V, Olkin I. Statistical methods for meta-analysis. Orlando, FL: Academic Press; 1985.

68. Kurtz ZD, Müller CL, Miraldi ER, Littman DR, Blaser MJ, Bonneau RA. Sparse and compositionally robust inference of microbial ecological networks. PLoS Comput Biol. 2015;11(5):e1004226.

69. Pedregosa F, Varoquaux G, Gramfort A, Michel V, Thirion B, Grisel O, et al. Scikit-learn: Machine Learning in Python. Journal of Machine Learning Research. 2011;12:2825–30.

70. Edge D, Trinh H, Newman Cheng JB, Chao A, Mody A, Truitt S, et al. From local to global: A graph rag approach to query-focused summarization, 2025. URL https://arxiv org/abs/240416130. 2025;2404.

71. Schoch CL, Ciufo S, Domrachev M, Hotton CL, Kannan S, Khovanskaya R, et al. NCBI Taxonomy: a comprehensive update on curation, resources and tools. Database (Oxford). 2020;2020.

72. Parks DH, Chuvochina M, Rinke C, Mussig AJ, Chaumeil PA, Hugenholtz P. GTDB: an ongoing census of bacterial and archaeal diversity through a phylogenetically consistent, rank normalized and complete genome-based taxonomy. Nucleic Acids Res. 2022;50(D1):D785–d94.

73. Agarwal S, Ahmad L, Ai J, Altman S, Applebaum A, Arbus E, et al. gpt-oss-120b & gpt-oss-20b model card. arXiv preprint arXiv:250810925. 2025.

74. Balachandran A. MedEmbed: Medical-Focused Embedding Models. 2024.

