## Supplementary figures and images for "Gut community-level analysis reveals an altered balance between *Phocaeicola vulgatus* and *Bacteroides fragilis* in Alzheimer’s disease"

### Extended Data Figure 1

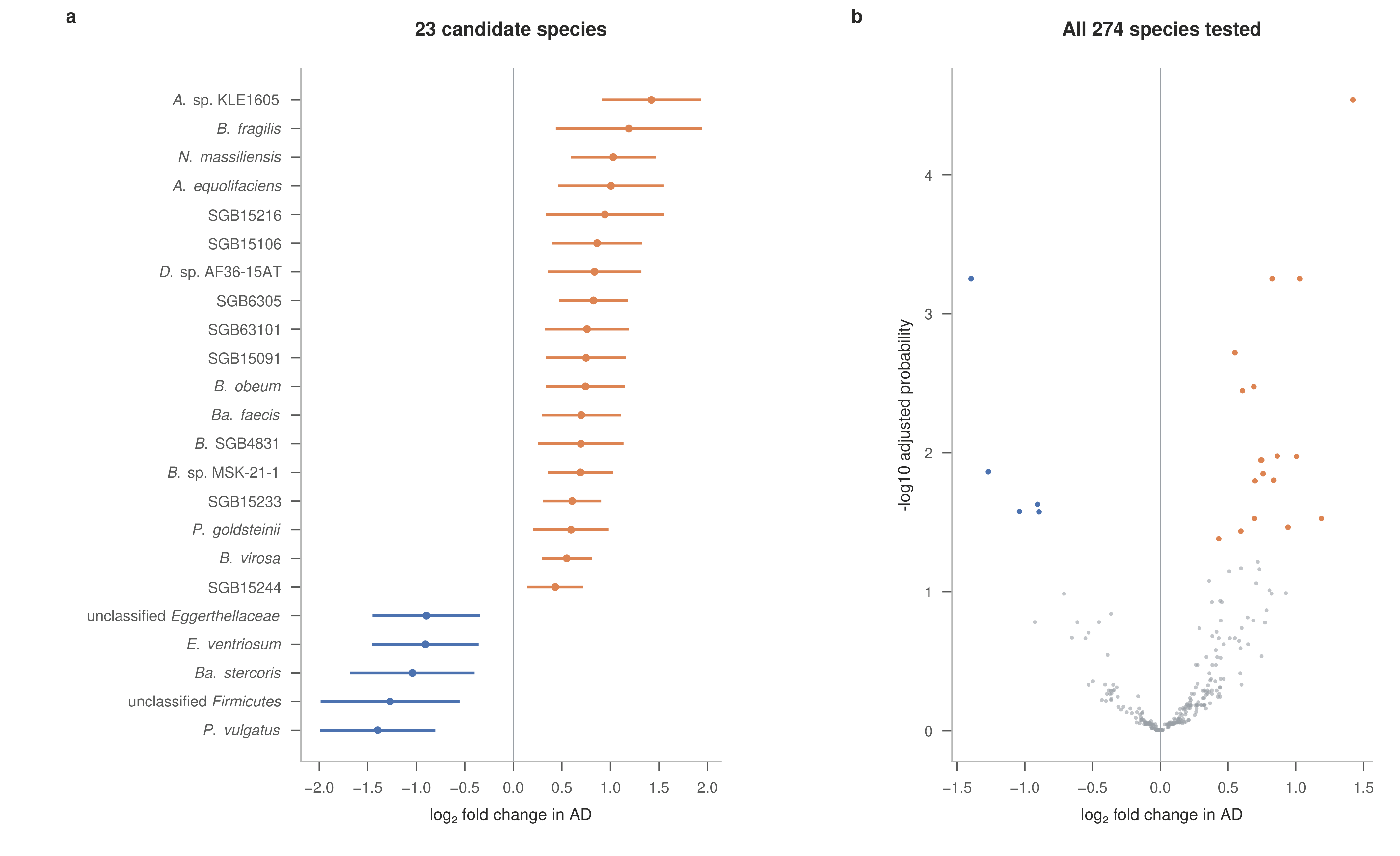

### Extended Data Figure 2

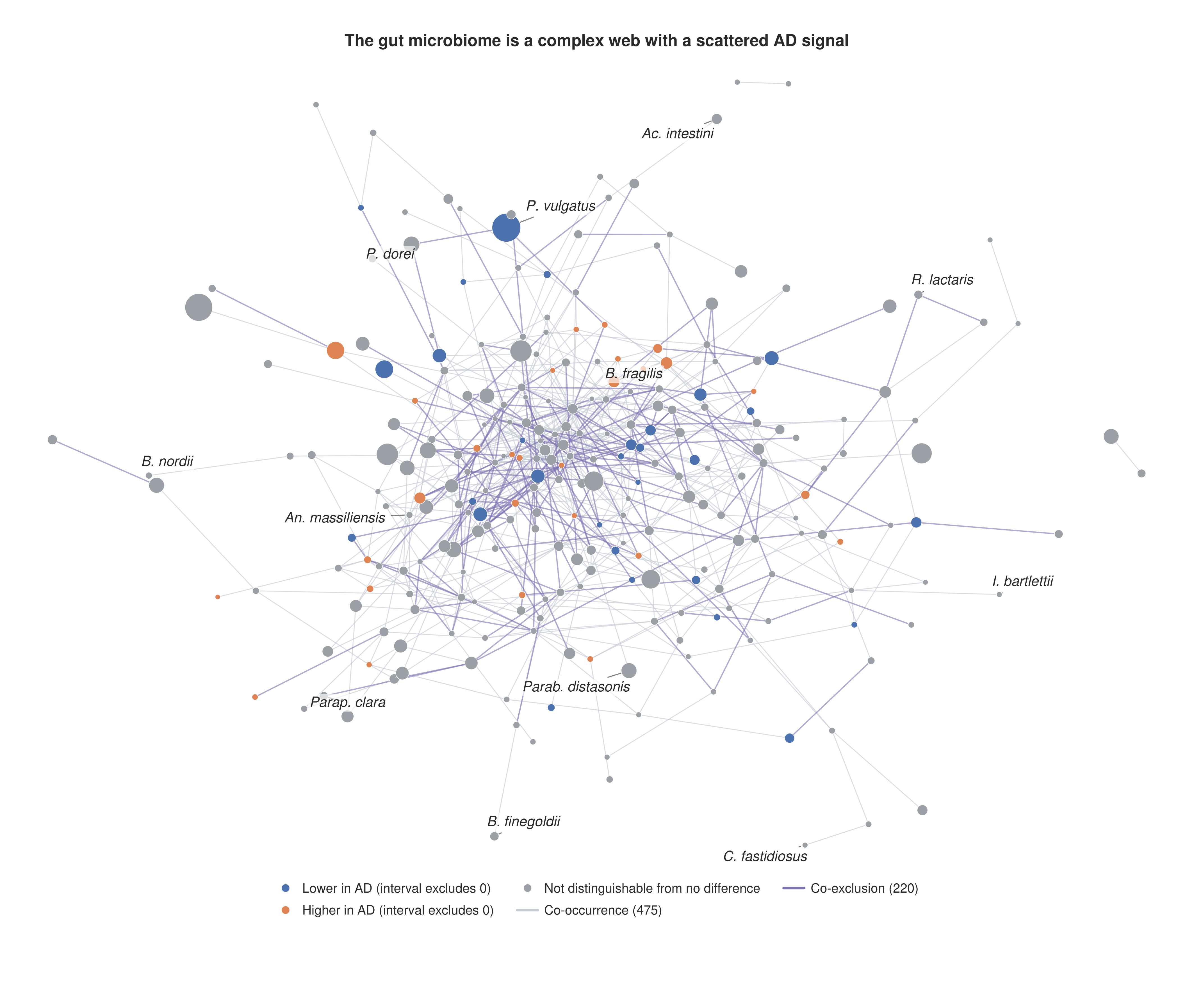

### Extended Data Figure 3

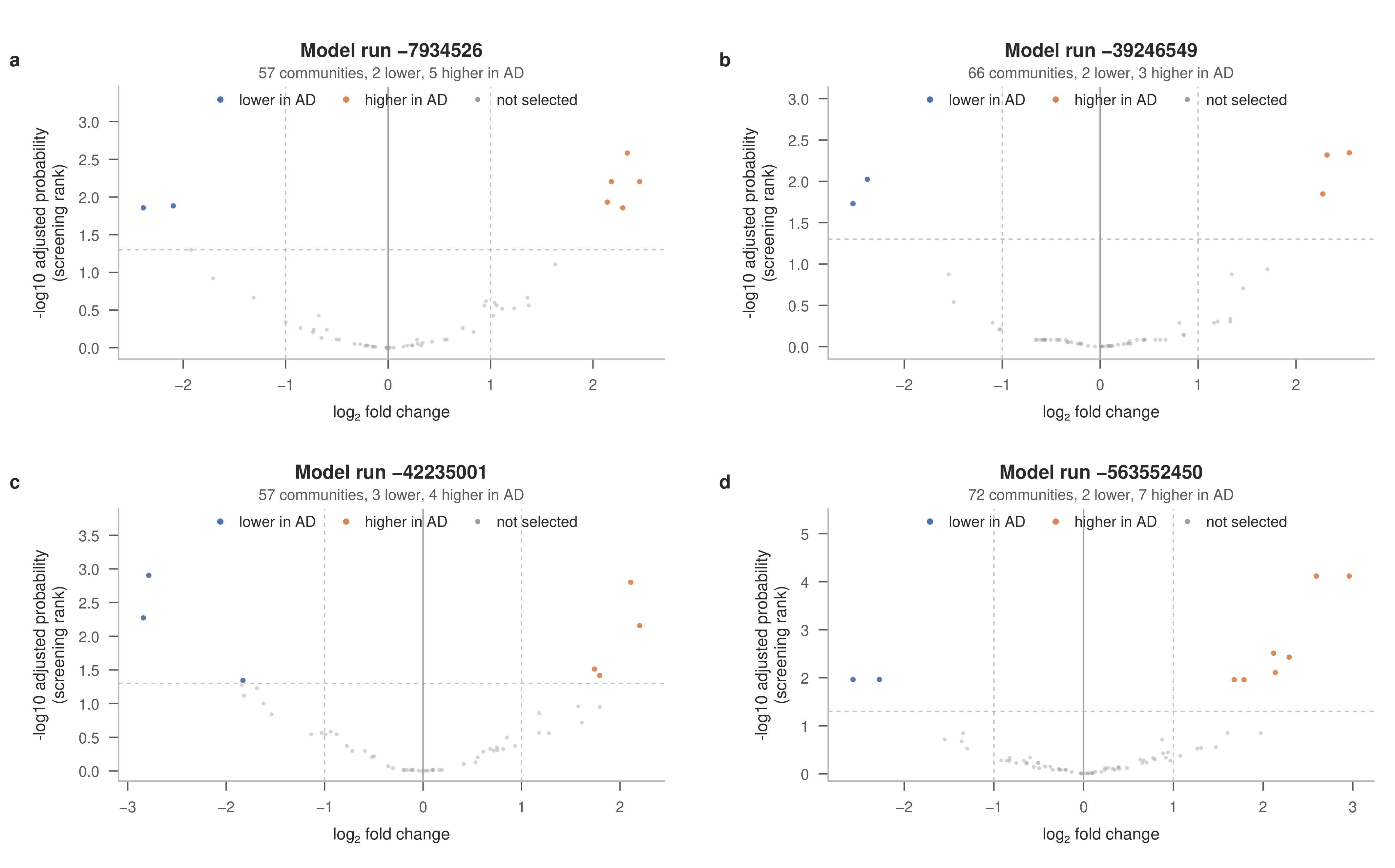

### Extended Data Figure 4

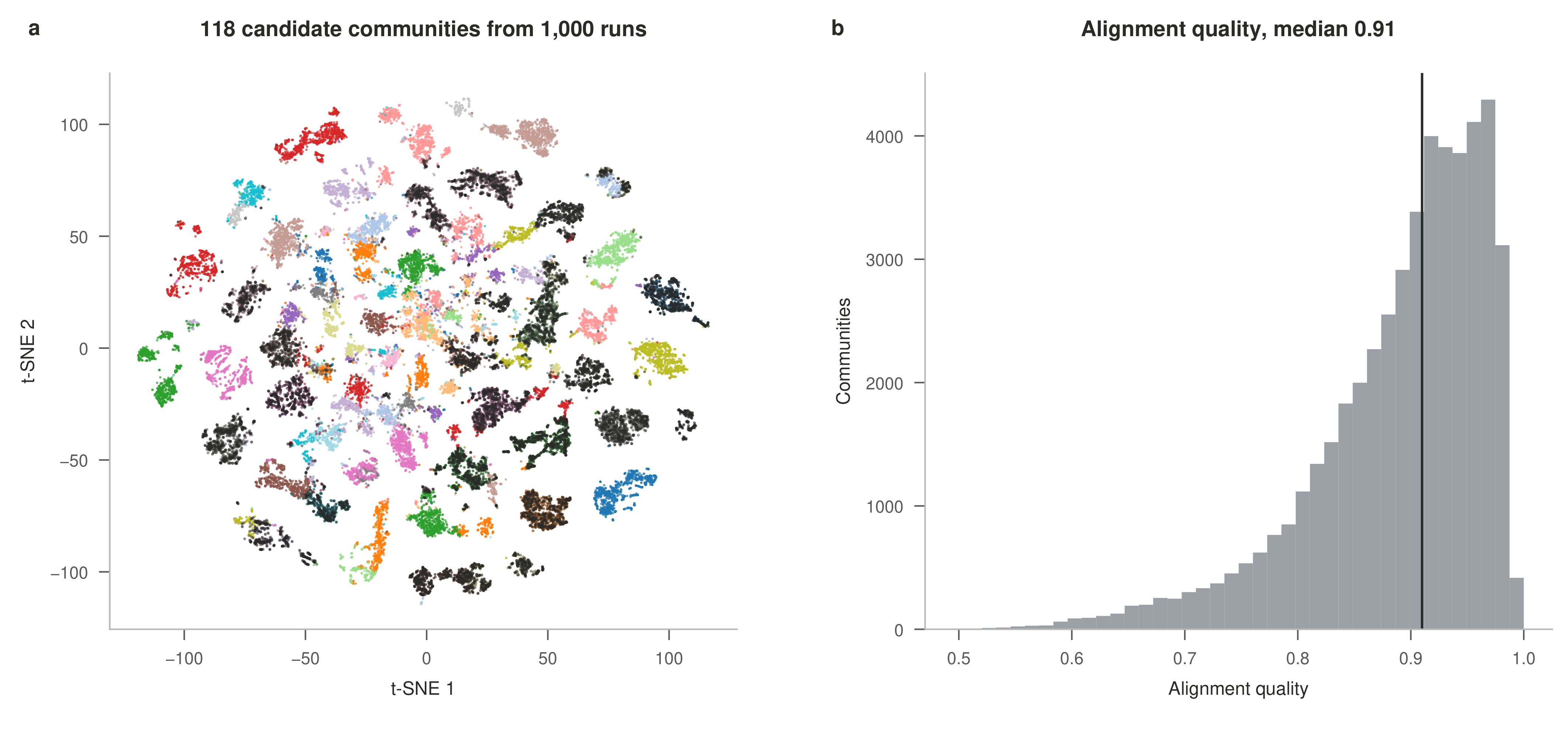

### Extended Data Figure 5

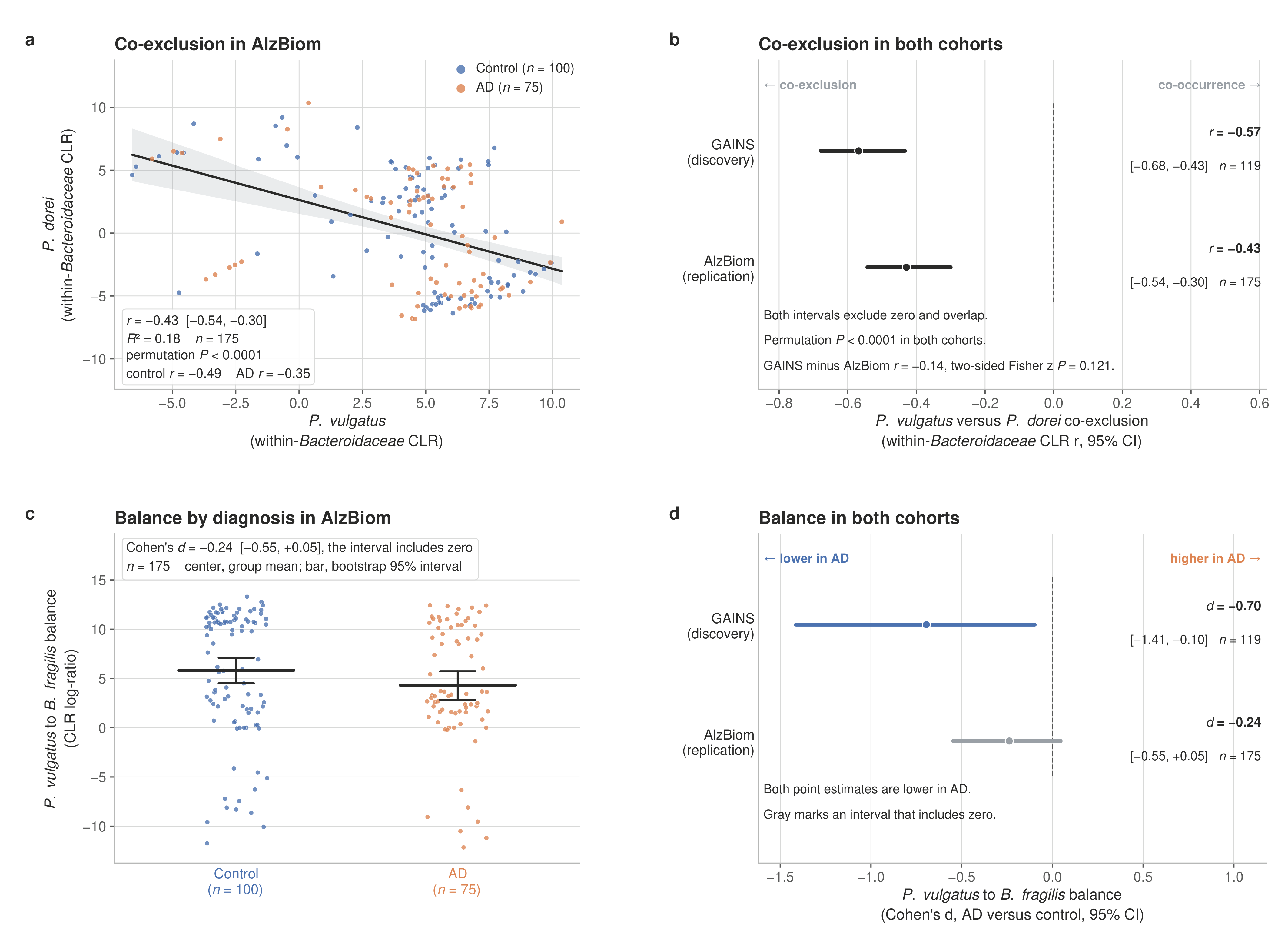

### Extended Data Figure 6

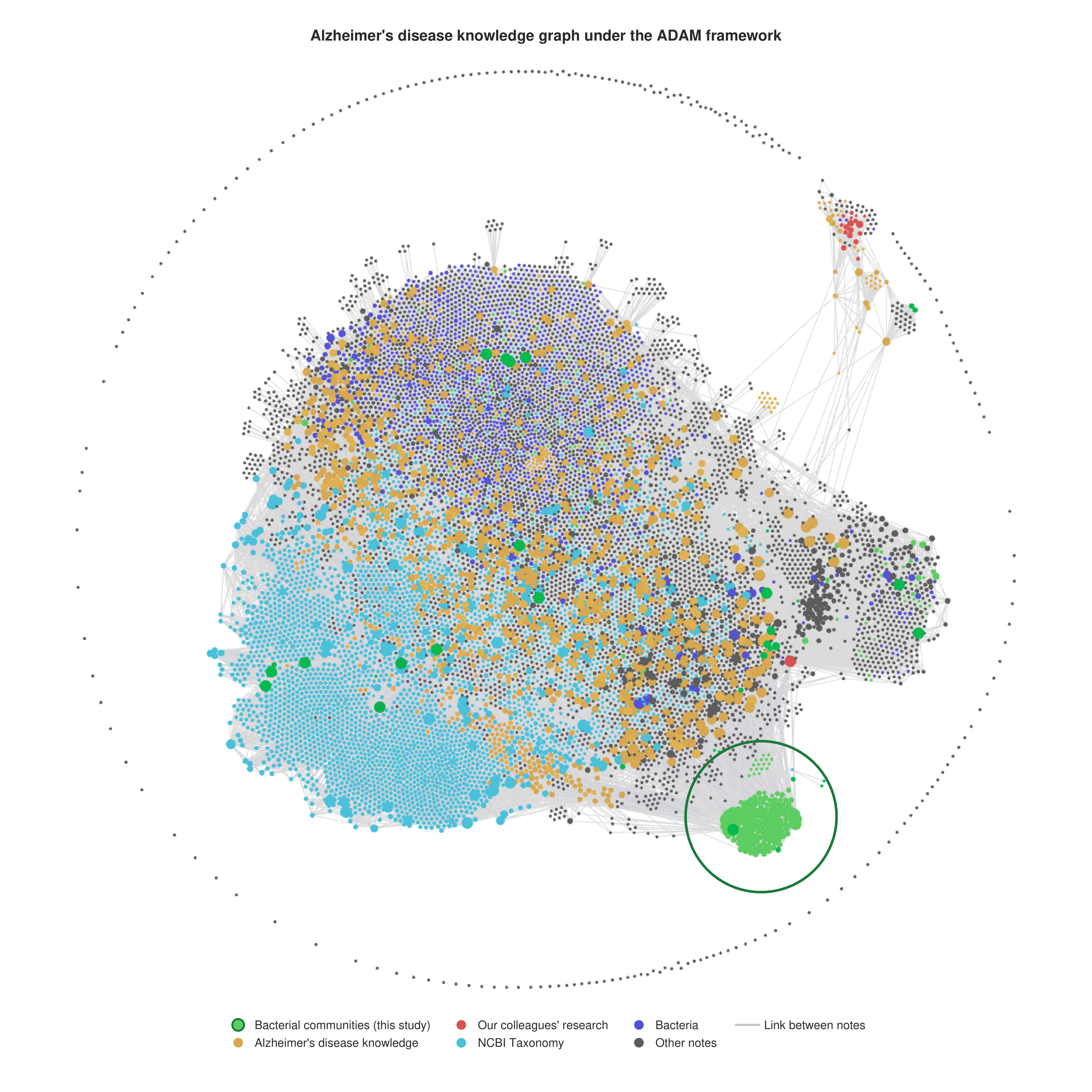

### Extended Data Figure 8

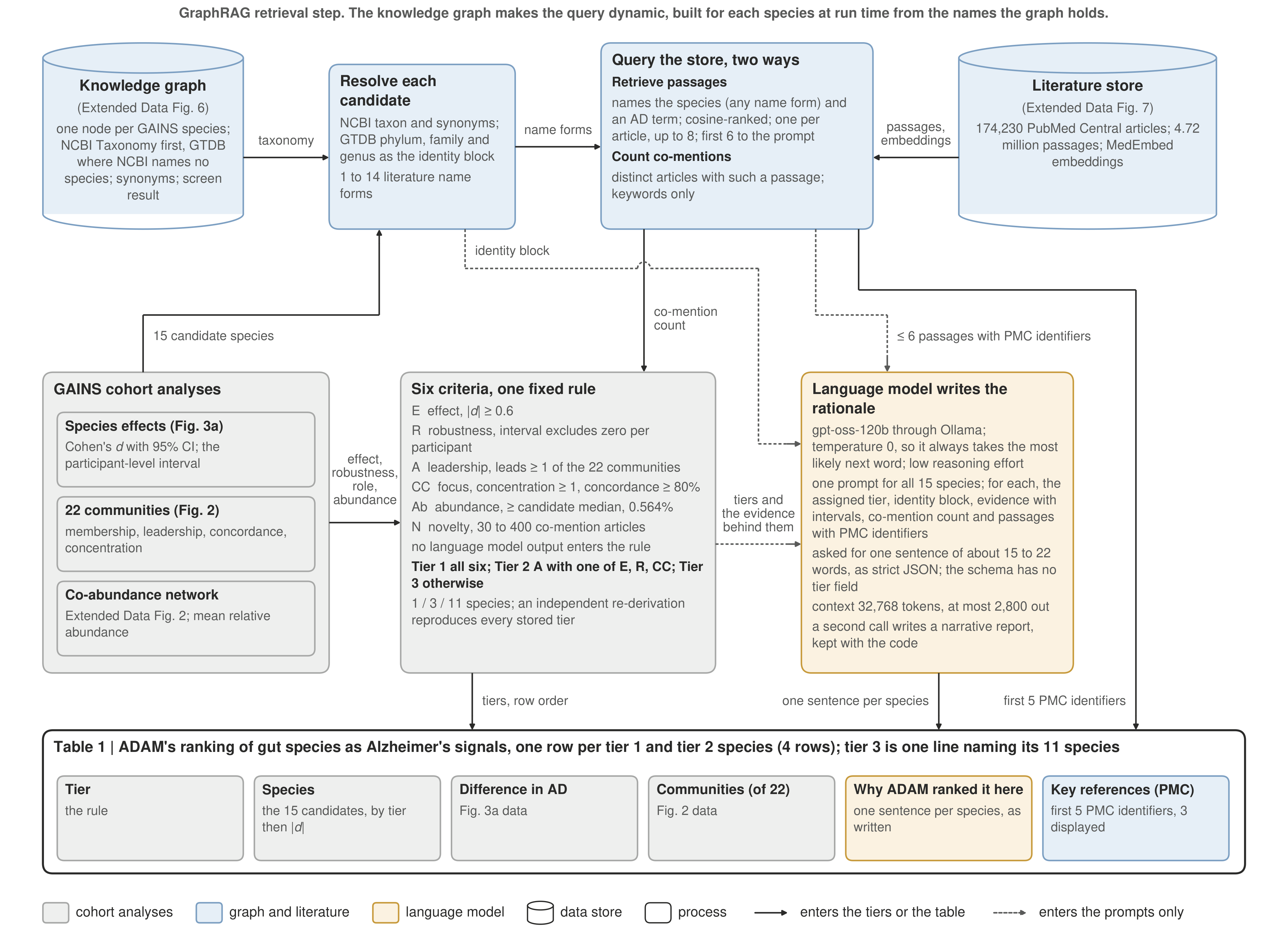

### Extended Data Table 1

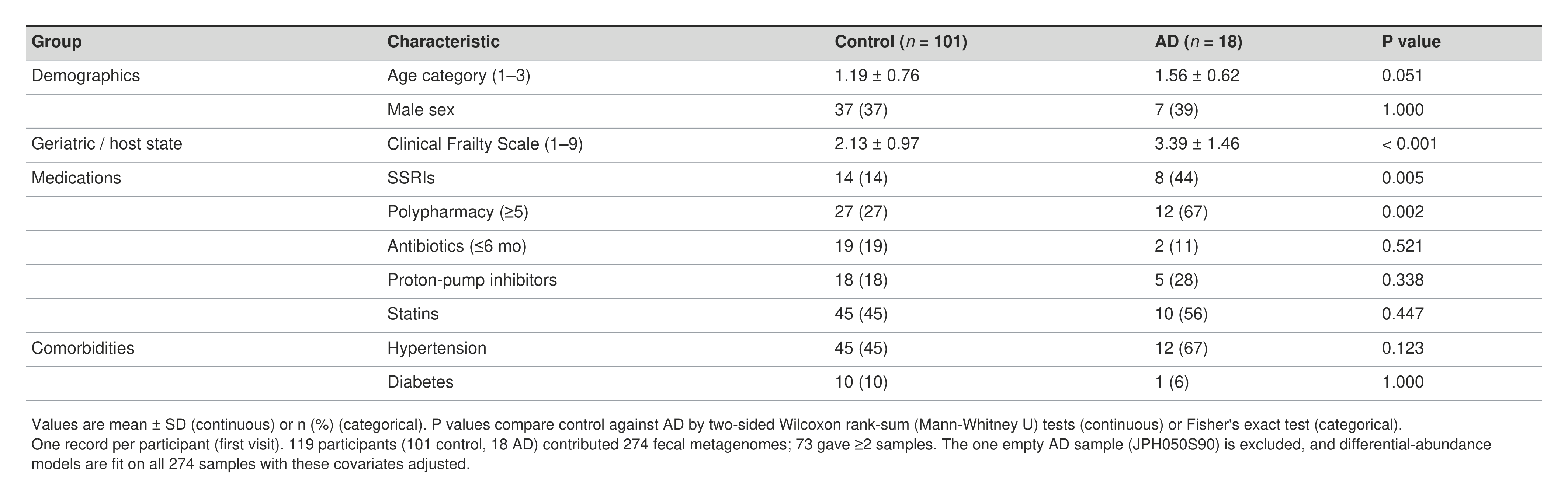

### Extended Data Table 2

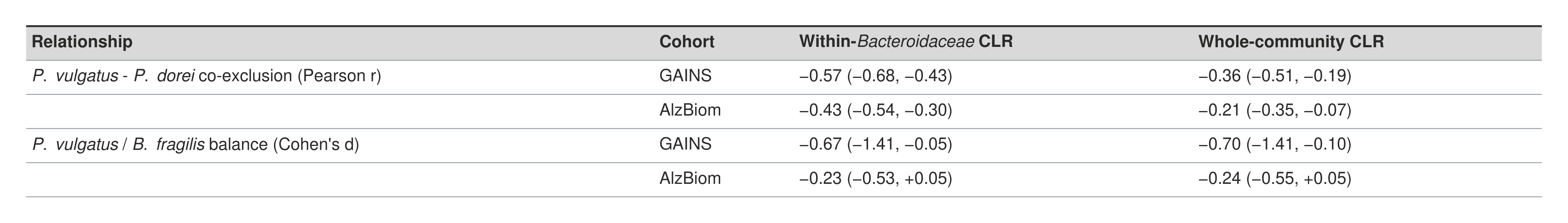
